# Single-nucleus profiling identifies accelerated oligodendrocyte precursor cell senescence in a mouse model of Down Syndrome

**DOI:** 10.1101/2023.04.17.537139

**Authors:** Bianca Rusu, Bharti Kukreja, Taiyi Wu, Sophie J. Dan, Min Yi Feng, Brian T. Kalish

## Abstract

Down Syndrome (DS), the most common genetic cause of intellectual disability, is associated with lifelong cognitive disability. However, the mechanisms by which triplication of human chromosome 21 genes drive neuroinflammation and cognitive dysfunction are poorly understood. Here, using the Ts65Dn mouse model of DS, we performed an integrated single-nucleus RNA and ATAC-seq analysis of the cortex. We identify cell type-specific transcriptional and chromatin-associated changes in the Ts65Dn cortex, including regulators of neuroinflammation, transcription and translation, myelination, and mitochondrial function. We discover enrichment of a senescence-associated transcriptional signature in Ts65Dn oligodendrocyte precursor cells (OPCs) and epigenetic changes consistent with a loss of heterochromatin. We find that senescence is restricted to a subset of cortical OPCs concentrated in deep cortical layers. Treatment of Ts65Dn mice with a senescence-reducing flavonoid rescues cortical OPC proliferation, restores microglial homeostasis, and improves contextual fear memory. Together, these findings suggest that cortical OPC senescence may be an important driver of neuropathology in DS.

## Introduction

Down syndrome (DS) is the leading genetic cause of intellectual disability worldwide, occurring in 1 in ∼800 live births^1, 2^. DS is caused by the presence of an extra copy or major portion of human chromosome 21 (HSA21)^3^. DS results in learning, memory, and language impairment, leading to lifelong cognitive disability, as well as a near-universal risk of early-onset neurodegeneration^4^. Triplication of HSA21 results in increased expression of ∼160 protein coding genes, as well as >300 genes of unknown coding or functional potential^5^. Overexpression of these genes results in complex perturbations of multiple processes involved in neurological development and function^6–10^.

DS is a unique opportunity for studying changes in brain aging across the lifespan, as DS individuals age prematurely and atypically^3^. Importantly, gene dosage imbalance of this relatively small set of genes has a profound cascading effect, including secondary changes in expression of hundreds of other genes, leading to disruption of several pathways important for neurogenesis, cell differentiation, synapse formation and plasticity, axon guidance, immune regulation, and myelination^6^. By 40 years of age, there is a ubiquitous occurrence of plaques and neurofibrillary tangles of hyperphosphorylated tau suggestive of Alzheimer’s Disease (AD), as well as clinical signs of dementia, including changes in sociability, language, and depressive symptoms^11–13^. Neuroinflammation – a key driver of neurodegeneration – is a neuropathologic hallmark of DS^14, 15^ and is in part driven by triplication of several immune-related genes, including four of the six interferon (IFN) receptors^16, 17^. There is also triplication of the amyloid precursor protein (App) and S100β, with the resultant overexpression of the pluripotent neuroinflammatory cytokine interleukin-1 (IL-1)^13, 18, 19^. Importantly, there is evidence of microglial activation in DS humans and mice with trisomy, and reducing over-activated microglia restores cognitive performance in trisomic mice^20^. While neuroinflammation is recognized as a core feature of DS, the molecular mechanisms and age-related progression of inflammation-associated aging (“inflammaging”), as well as the relationship to cognitive decline, are poorly understood.

To address these gaps, we leveraged single-nucleus sequencing to identify candidate pathways perturbed in the brain of a well-characterized DS mouse model. Ts65Dn mice contain a partial triplication of chromosome 16 (MMU16) corresponding to trisomy of ∼55% of HSA21-coding orthologs^21–23^. These trisomic mice recapitulate many of the characteristics of human DS, including learning, memory, and cognitive deficits, as well as developmental delays and oligodendrocyte (OL) maturation deficits^21, 23, 24^. Through multi-omic profiling of the Ts65Dn brain, we identify broad disruptions in pathways associated with neuroinflammation, transcriptional and translational regulation, mitochondrial and ribosomal dysfunction, and the integrated stress response (ISR) at both the epigenomic and transcriptomic level across neuronal and non-neuronal cell populations. We further identify a senescence-associated gene signature in Ts65Dn oligodendrocyte precursor cells (OPCs). Using orthogonal approaches, we validate our sequencing findings and discover that a subset of cortical OPCs are prematurely senescent in the trisomic mouse brain. We show that treatment of Ts65Dn mice with the anti-senescence flavonoid, Fisetin, has restorative effects on cortical OPCs, including reducing senescence-associated-β-gal (SA-β-gal) activity, increasing proliferation and progenitor abundance, restoring microglia homeostasis, and improving contextual fear memory. In sum, this work identifies novel cellular mechanisms that drive chronic neuroinflammation and cognitive dysfunction in the Ts65Dn DS mouse model.

## RESULTS

### Characterization of cell types in the adult Ts65Dn cortex using multi-omic single nucleus sequencing

To identify cellular programs underlying age-related cognitive decline in trisomy, we performed single-nucleus RNA-seq (snRNA-seq) and ATAC-seq (snATAC-seq) on the 6-month (6mo) Ts65Dn and euploid control (CTL) littermate mouse cortex using the 10X Genomics platform (Figure 1A; Methods)^25, 26^. Each biological sample was split into two parts to capture and analyze both the mRNA and accessible chromatin: one part was used for snRNA-seq and the other was used for snATAC-seq. Ts65Dn mice and littermate CTLs were selected from three separate litters. The cerebral cortex, the outermost portion of the brain, was chosen as the region of interest for this study due to its involvement in cognition and higher-order processing^27^, and its known structural and function deficits in DS individuals, including reductions in cortical volume and surface area, as well as decreased cell counts, abnormal synapto-dendritic processes, and disorganized cortical lamination^28, 29^. Through multi-omic single-nucleus sequencing, we analyzed the transcriptomes and epigenomes from each dataset using the well-established Seurat^30^ and Signac^31^ pipelines, respectively. Briefly, data was quality control (QC) filtered to remove clusters likely to be of low quality resulting from debris, doublets, and dead cells, and was subsequently normalized using default parameters (Methods). After QC processing, we profiled a total of 37,251 nuclei for snRNA-seq and 53,244 nuclei for snATAC-seq (Figure 1–figure supplement 1A-B).

**Figure 1:**
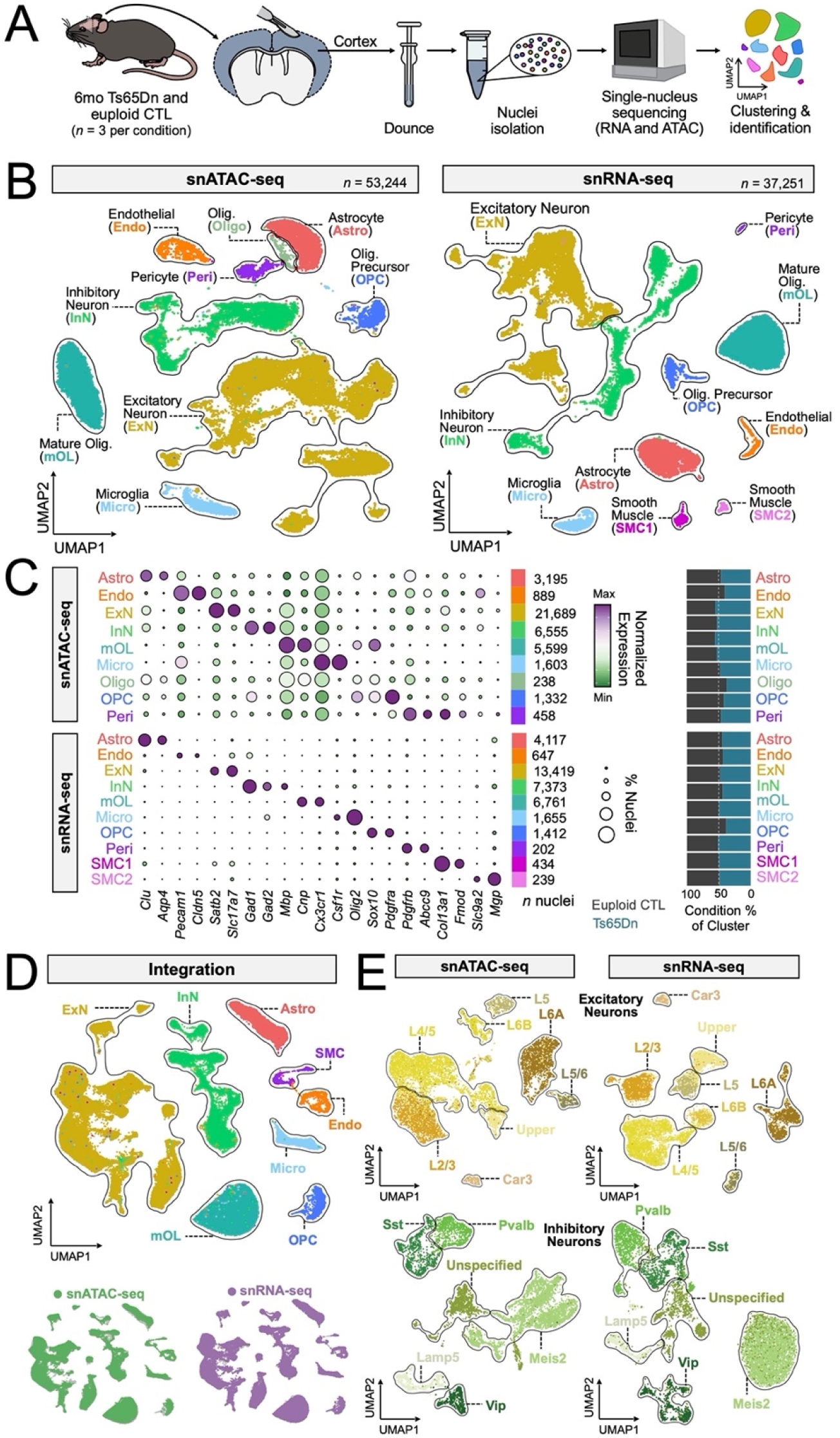
Multi-modal single-nucleus sequencing of the Ts65Dn mouse cortex. (A) Schematic representation of biological samples, cortical dissection, tissue processing and sequencing workflow for snATAC- and snRNA-seq from 6mo Ts65Dn and euploid control (CTL) mice. *n* = 3 biological replicates per condition. (B) UMAP visualizations of snATAC- and snRNA-seq datasets, where each dot represents a single nucleus, for a total of 53,244 and 37,251 nuclei profiled with snATAC- and snRNA-seq, respectively. UMAP plots are generated from combined replicates across Ts65Dn and CTL conditions. Each cluster is colored by cell type: Astro, astrocytes; Endo, endothelial cells; ExN, excitatory neurons; InN, inhibitory neurons; mOL, mature oligodendrocytes; Micro, microglia; Oligo, oligodendrocytes; OPC, oligodendrocyte precursor cells; Peri, pericytes; SMC1/2, smooth muscle cell subsets. (C) Dotplot of snATAC- and snRNA-seq datasets, showing gene expression or gene accessibility patterns, respectively, with several key canonical marker genes used for cluster identification. The diameter of the dot corresponds to the proportion of nuclei expressing or exhibiting accessibility of the indicated gene, and the color of the dot corresponds to the average expression or accessibility of the gene relative to all cell types. The number of nuclei assigned to each cell type are indicated. Barplots on the right depict the fraction of nuclei per cell type by condition. (D) UMAP visualization of multi-omic integration of snATAC- and snRNA-seq datasets colored by cell type assignment. On the right are UMAPs of the integration colored by originating dataset. (E) UMAP visualizations of excitatory and inhibitory neuron subsets as extracted from snATAC- and snRNA-seq datasets. Each cluster is colored by cell type. For excitatory neuron UMAPs: Car3, Car3 expressing excitatory neurons; L2/3, cortical layer II-III excitatory neurons; L4/5, cortical layer IV-V excitatory neurons; L5, cortical layer V excitatory neurons; L5/6, cortical layer V-VI excitatory neurons; L6A/B, cortical layer VI excitatory neuron subsets; Upper, mixed upper layer (II-IV) excitatory neurons. For inhibitory neuron UMAPs: Lamp5, Lamp5-expressing interneurons; Meis2, Meis2-expressing interneurons; Pvalb, parvalbumin-expressing interneurons; Sst, somatostatin-expressing interneurons; Vip, vasoactive intestinal peptide-expressing interneurons; Unspecified, interneurons of unspecified classification.

Genes with high variance were used to compute principal components for projecting and clustering cell populations with similar molecular signatures. Unsupervised clustering through uniform manifold approximation and projection (UMAP; Methods) dimensionality reduction and Leiden clustering revealed 10 major cell classes across the snRNA-seq and snATAC-seq datasets: astrocytes (Astro), excitatory neurons (ExN), inhibitory neurons (InN), mature oligodendrocytes (mOL), microglia (Micro), oligodendrocyte precursor cells (OPC), oligodendrocyte lineage cells (Oligo), endothelial cells (Endo), pericytes (Peri), and smooth muscle cells (SMC) (Figure 1B). Cell types were assigned based on a combination of canonical marker gene expression (Figure 1C) and confirmed through a machine learning reference-based mapping approach (scPred; Methods)^32^ using the Allen Brain Atlas whole cortex and hippocampus 10X Genomics dataset (Methods)^33^. All major cell types were present in both conditions, with high transcriptomic and epigenetic overlap between genotypes. Label transfer and integration of snRNA-seq to snATAC-seq further confirmed cell population assignments and demonstrated strong concordance between the two modalities, with cell types identified in either platform grouping together in the integrated UMAP space (Figure 1D; Methods).

The ExN and InN clusters were proportionally the largest by number out of all identified cell type clusters, consistent with previous reports that neurons are the most numerous cell type in the cortex^34^. Thus, in both snATAC-seq and snRNA-seq, we performed a second level of hierarchical clustering on ExNs and InNs to identify distinct sub-populations of each neuron class (Figure 1E). Again, using the scPred algorithm on the snRNA-seq dataset, we identified clusters spanning 5 cortical layers (L2-6), a distinct Car3+ population of ExNs, as well as 5 sub-populations of InNs, all with distinct transcriptional signatures. Integration of snRNA-seq and snATAC-seq neuronal sub-clusters revealed corresponding ExN cortical layer and InN subpopulations within the snATAC-seq dataset (Figures 1–figure supplement 1C-D). Overall, this analysis transcriptionally and epigenetically identified all major neuronal and non-neuronal cell types and subtypes across the murine cortex.

### Trisomy-associated transcriptomic changes are highly cell type specific

To identify transcriptional signatures of trisomy-associated cognitive decline, we performed cell type-specific differential gene expression (DGE) between Ts65Dn and euploid CTL (Methods). We used differentially expressed genes (DEGs) that met a false discovery rate (FDR) value of 5% to define statistically significant changes in transcript-level expression between Ts65Dn and CTL offspring. We identified cell type-specific DEGs in all cell populations (Figure 2A), with 64.3% of DEGs upregulated (log_2_FC > 0) and 35.7% downregulated (log_2_FC < 0) in Ts65Dn mice. We found that 38% of the DEGs significantly altered by Ts65Dn trisomy were cell type-specific, with the remaining 62% intersecting between two or more cell types (Figure 2B).

**Figure 2:**
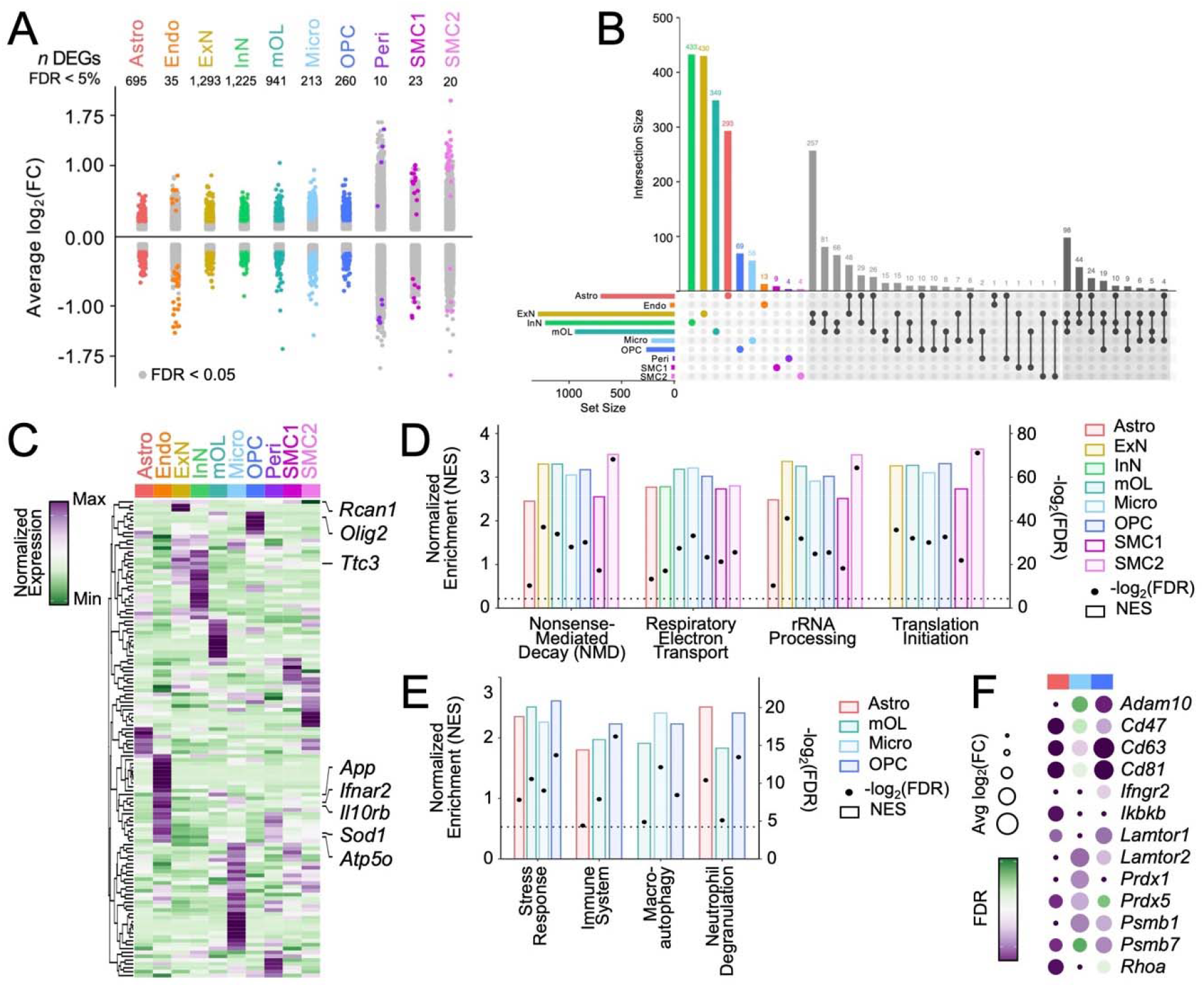
Transcriptionally distinct cell subpopulations in Ts65Dn cortex. (A) Strip plot displaying differentially expressed genes (DEGs) between Ts65Dn and CTL offspring at 6mo. Colored dots represent significant genes (FDR < 5%) per cell type. The number of significant DEGs per cell type is indicated. The *x*-axis displays all major cortical cell types profiled through snRNA-seq. Data obtained from *n* = 3 biological replicates per condition. (B) UpSet plot displaying the number of unique and shared DEGs across cell types, with unique genes colored based on cell type, and genes shared between 2 or 3 cell types indicated by black dots connected by lines according to shared origins. The histogram indicates the number of DEGs for each cell type, and the barplots show the number of significant DEGs (FDR < 5%) per cell type. (C) Hierarchically clustered heatmap of normalized gene expression for genes contained within the Ts65Dn chromosomal product. The plot displays gene activity across al cell types in snRNA-seq. Several genes of interest are indicated. Each column represents a cell type. The color code represents the row-normalized expression for each gene. (D) Gene set enrichment analysis (GSEA) of several Reactome biological pathways enriched across a majority of cortical cell types. The bars correspond to the left *y*-axis, displaying normalized enrichment score for each cell type by pathway and are colored by cell type identity. The dots correspond to the right *y*-axis, displaying -log_2_(FDR) for each cell type by pathway; the dashed horizontal line intercepts the right *y-*axis at 4.3, corresponding to an FDR = 0.05 (5%). All dots above this horizontal line are statistically significant. (E) as in (D), but depicting GSEA of several Reactome biological pathways enriched across astrocytes (Astro), mature oligodendrocytes (mOL), microglia (Micro), or oligodendrocyte precursor cells (OPC). (F) Dotplot of select differentially expressed genes (DEGs) within astrocytes (red), microglia (light blue), and OPCs (dark blue). Each row represents a gene. The color code represents the FDR, and the size of the dots represents the average log_2_fold change (avg log_2_(FC)) of gene expression between Ts65Dn and CTL.

The highest number of shared DEGs was found between ExNs and InNs, reflecting their close biological function and identity (Figure 2B).Several genes were found to be overexpressed in a majority of ExN sub-clusters (at least 5 out of 8), including *Calm1*, an essential regulator of early neuronal migration^35^, *Fth1*, a ferroxidase enzyme that supports iron detoxification as part of the neuronal antioxidant defense system^36^, *Scg5*, a secretory chaperone that co-localizes with aggregated proteins in neurodegenerative diseases^37^, and *Pcp4*, a modulator of calcium signaling that is triplicated in DS^38^. A similar trend was observed in InNs, with several genes showing overexpression across multiple subtype clusters (at least 4 out of 6), including the aforementioned *Fth1* and *Scg5*, as well as *Hspa8*, a molecular chaperone whose expression increases during injury and activates pro-inflammatory responses through NF-κB signaling^39^. Several large (*Rpl*) and small (*Rps*) ribosomal subunit genes, including *Rpl19/26/27a* and *Rps7* were also overexpressed in several InN subtypes, suggesting a disruption in translation-associated ribosomal machinery.

All cell types exhibited upregulation of ∼10-20% of the genes expressed on the Ts65Dn trans chromosome at the transcript level. However, not all Ts65Dn triplicated genes were equally altered in each cell type (Figure 2C). Only a few genes such as *Atp5o* or *Sod1* were consistently found at higher levels in trisomic mice across various cell types, while others were overexpressed only in particular cell populations. For instance, *Olig2*, a key transcription factor (TF) that activates the expression of myelin associated genes in OL-lineage cells^40, 41^, was upregulated in trisomic OPCs by 15%; its overexpression is known to contribute to impaired OPC proliferation and differentiation in DS^7, 8, 42^. *Ttc3*, an interactor of nerve growth factor (Ngf) that strongly inhibits neurite extension upon overexpression^43^, was upregulated in Ts65Dn ExNs by 13%. *App*, associated with increased amyloid-β (Aβ) aggregation and plaque deposition^44^ showed enrichment of 8.6% and 11% in ExNs and microglia, respectively, and *Rcan1*, whose overexpression leads to neurofibrillary tangles^45^, was enriched in Ts65Dn endothelial cells by 16%. Lastly, pericytes and astrocytes exhibited an increase in *Ifnar2* expression by 37% and 8.6%, respectively, while microglia showed an increased expression of *Ifnar2* by 10% and *Il10rb* by 13%; together, these interferon (IFN) receptors may contribute to the hyper-inflammatory milieu of the trisomic brain.

### Gene set enrichment analysis reveals dysregulation of several biological pathways in Ts65Dn mice

To identify functional pathways perturbed in Ts65Dn mice, we performed gene set enrichment analysis (GSEA) using the Reactome repertoire of biological pathways^46^ on DEGs across all cell types (Methods). Of the top 10 enriched GSEA categories, 4 emerged as common processes altered in almost all cell types: ribosomal RNA (rRNA) processing, respiratory electron transport, nonsense mediated decay (NMD), and translation initiation (Figure 2D). DS is characterized by an impairment in mitochondrial and ribosomal biogenesis, as well as integrated stress response (ISR)-mediated disruption in proteostasis^47–49^. Consistent with this finding, we observed overexpression and dysregulation of over 30 large (*Rpl*) and small (*Rps*) ribosomal subunit genes and 15 mitochondrial (*mt* and *cox*) transcripts in the Ts65Dn brain. Genes involved in translation initiation, such as *eIF4e, eIF3e*, and *eIF3a*^50, 51^, were also found to have increased expression in all neuronal and non-neuronal cortical Ts65Dn cells. DEGs in trisomic astrocytes, microglia, and OPCs also exhibited enrichment of GSEA categories including immune activation, neutrophil degranulation, macroautophagy, and stress response (Figure 2E). In Ts65Dn astrocytes, we observed upregulation of *Ikbkb*, an activator of the NF κB pathway^52^, *RhoA*, a regulator of reactive astrocyte dynamics^53^, and inflammatory mediators *Cd47*, *Cd63*, and *Cd81*^54^ (Figure 2F). Ts65Dn OPCs likewise exhibited increased expression of several genes whose functions have been associated with immune response, such as *Adam10*, an extracellular matrix (ECM) protein involved in inflammatory signaling^55^ and *Ifngr2*, an IFN receptor, whose upregulation induces increased immunomodulatory signaling^56^. Trisomic astrocytes, microglia, and OPCs also showed increased expression of several proteasome subunits including *Psmb1/7*^57^, as well as antioxidant enzymes *Prdx1/5*, whose functions are important during periods of oxidative stress^58^, and *Lamtor1/2*, which mediate cellular autophagy and survival responses to inflammation^59^. Taken together, these transcriptional changes are consistent with previous reports of glial reactivity in the Ts65Dn cortex^20, 60–62^.

### Cortical Ts65Dn OPCs exhibit enrichment for a senescence-associated gene signature

Given reports of increased DNA damage and chronic neuroinflammation in DS and the Ts65Dn brain^63–65^, we hypothesized that subpopulations of Ts65Dn cells would exhibit accelerated senescence. Senescent cells accumulate in aged tissues due to exhaustion of proliferation-competent cells^66, 67^. However, the identification and characterization of senescent cells, particularly in bulk or single-nucleus sequencing data, is challenging due to the low average expression of many senescence-associated transcripts and the the imprecise genetic definition of cellular senescence. We thus performed GSEA using a recently-published senescence-associated gene signature, SenMayo, which is composed of 125 senescence-associated genes validated on murine and human age-related datasets^68^ (Methods). Through this analysis, we observed selective enrichment (normalized enrichment score, NES > 0, FDR < 5%) of the SenMayo gene set only in Ts65Dn OPCs (Figures 3A and 3B), suggesting a cell type specific senescence phenotype. Trisomic OPCs further demonstrated altered expression of several genes pertaining to maturation, proliferation, inflammation, and senescence (Figure 3C). We found increased expression of *Gpr17*, which causes myelinogenesis defects when overexpressed in OL lineage cells^69, 70^, *Mif*, a pro-inflammatory cytokine that regulates NF-κB and p53 expression^71, 72^, as well as *Mmp15* and *Adamts17*, which are endoproteases that mediate the activity and bioavailability of inflammatory factors, such as TNFα and IL-1^73, 74^. GSEA analysis in Ts65Dn OPCs also revealed an enrichment for metabolic regulation by p53, a key transcription factor (TF) whose upregulation induces growth arrest, apoptosis, and cellular senescence^75^.

**Figure 3:**
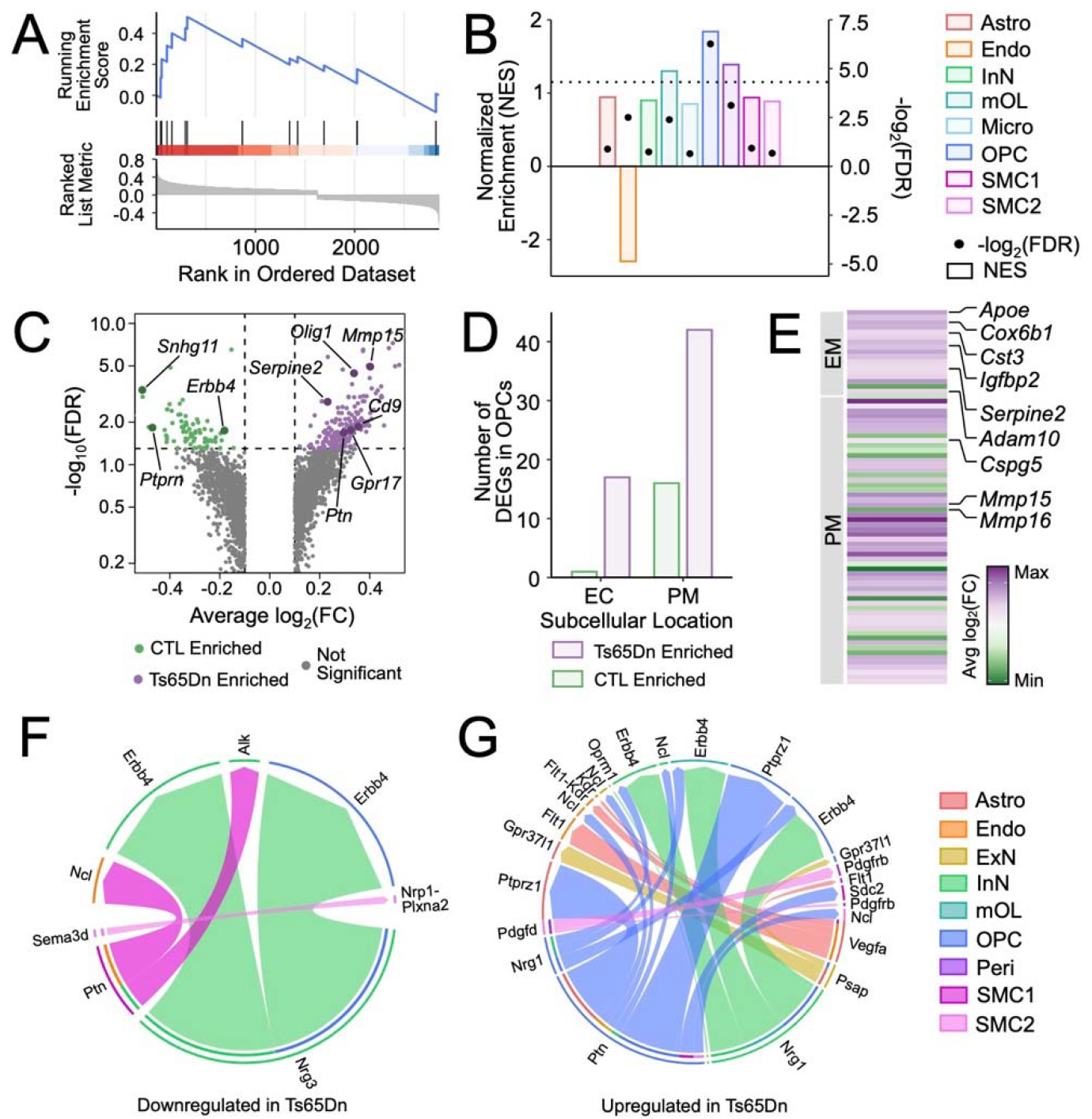
Cortical Ts65Dn OPCs exhibit a selective senescence-associated phenotype. (A) GSEA-based enrichment plot of the SenMayo senescence gene signature for OPCs. The upper *y*- axis represents the enrichment score (ES), and the blue line represents the running enrichment score. The *x*-axis displays gene ranked according to their expression in Ts65Dn, with the most upregulated genes on the left-hand side and the most downregulated genes towards the right-hand side. Black vertical lines depict the positions of individual genes and their enrichment within the transcriptional signature. (B) Barplot displaying the enrichment of the SenMayo senescence gene signature across cell types. The bars correspond to the left *y*-axis, displaying normalized enrichment score for each cell type by pathway and are colored by cell type identity. The dots correspond to the right *y*-axis, displaying - log_2_(FDR) for each cell type by pathway; the dashed horizontal line intercepts the right *y-*axis at 4.3, corresponding to an FDR = 0.05 (5%). All dots above this horizontal line are statistically significant. (C) Volcano plot of DEGs in Ts65Dn versus CTL conditions. Only genes with an average log_2_fold change (avg log_2_(FC)) > 0.1 or < -0.1. are included in the plot. Each dot represents a gene. Dots are colored according to enrichment: green dots are enriched in CTL, purple dots are enriched in Ts65Dn, and grey dots are not significantly enriched in either condition. The horizontal line depicts an FDR < 5%, such that all genes above this line are statistically significant. (D) Barplot depicting the number of DEGs (FDR < 5%) in OPCs with functionality localized to the extracellular compartment (EC) or plasma membrane (PM). Purple bars depict DEGs enriched in Ts65Dn, and green bars depict DEGs enriched in CTL. (E) Heatmap of differentially expressed genes (DEGs) localized to the extracellular space (EC) or plasma membrane (PM) in OPCs. Each row represents a gene. The color code represents the avg log_2_(FC) of gene expression between Ts65Dn and CTL. (F) Chord diagram of cell type signaling pathways that are downregulated in Ts65Dn versus CTL. Cell type identity of the ligand is indicated in the outermost edge of the diagram, while the cell identity of the receptor is indicated by the internal ring. Colored arrows indicate the specific ligand/receptor pairs and are colored according to the outgoing ligand signal. (G) as in (F) but depicting signaling pathways that are upregulated in Ts65Dn versus CTL.

Senescence is also marked by the widescale disruption of protein secretion and expression, which leads to the acquisition of a senescence-associated secretory phenotype (SASP). The SASP includes chemokines, cytokines, matrix metalloproteinases (MMPs), interleukins, and proteases, which together damage and modify the surrounding microenvironment^76^. Further examination of Ts65Dn OPC DEGs revealed that upregulated DEGs include several SASP family members, including *Apoe*, *Igfbp2*, *Cst3*, *Serpine2*, and *Cox6b1* (Figures 3D and 3E). Aberrant apolipoprotein E (*ApoE*) accumulation has been shown to drive senescence through degradation of nuclear envelope proteins that lead to heterochromatin destabilization and disorganization^77^. *Igfbp2* similarly induces and sustains cellular senescence by inhibiting apoptosis through the interaction with and protection of p21^78^. Alongside the induction of tissue sensing programs and secreted mediators, senescent cells demonstrate remodeling of the cell-surfaceome landscape through the upregulation of transcripts encoding plasma membrane (PM) proteins^79^. Indeed, many of the upregulated DEGs in Ts65Dn OPCs encoded PM proteins, such as ECM molecules (*Adam10*, *Cspg5*, *Mmp15/16*), and VEGF signaling molecules (*Cadm4*, *Cd63*) (Figures 3D and 3E). Taken together, the increase in extracellular (EC) and PM-encoding transcripts suggests an enhanced capability to sense and secrete environmental cues permitting senescent Ts65Dn OPCs an increased interaction with and influence of their surrounding microenvironment.

### Ligand-receptor analysis of the Ts65Dn cortex identifies changes in intercellular communication

Given the importance of cell-cell signaling in shaping neural circuits, we performed ligand receptor (LR) mapping in the Ts65Dn and euploid CTL brain using CellChat^80^ (Methods). We sought to build a comprehensive intercellular signaling network by leveraging the transcriptional profiles of each cell population to better understand how trisomy-driven changes in gene expression affect cell-cell signaling. Network analysis showed that 24 signaling pathways in the cortex were perturbed in Ts65Dn mice: pleiotrophin (*Ptn*) and neuregulin (*Nrg*) were amongst the top mediators of disrupted crosstalk between various major cell types in the Ts65Dn cortex (Figures 3F and 3G).

*Ptn* is a ligand for several receptors in the brain, including protein tyrosine phosphatase receptor type Z1 (*Ptprz1*), anaplastic lymphoma kinase (*Alk*), and nucleolin (*Ncl*)^81, 82^. We found increased *Ptn* to *Ptprz1* signaling in OPCs, decreased *Ptn* to *Alk* signaling between SMC1 and InNs, as well as decreased *Ptn* to *Ncl* signaling between SMC1 and endothelial cells in the trisomic mouse brain (Figures 3F and 3G). Upregulated *Ptn*-*Ptprz1* signaling indirectly increases the bioavailability of β-catenin, whose activity can hinder both developmental myelination and adult remyelination^83, 84^, while *Ptn*-*Alk* signaling promotes differentiation, growth, and survival^85^. Lastly, *Ncl* is expressed in endothelial cells and plays a role in angiogenesis and vascularization^86^. The downregulated signaling of a pro angiogenic LR interaction in endothelial cells is thus consistent with prior observations of impaired endothelial vascular recruitment and mobilization in DS^86, 87^.

Another set of perturbed pathways detected in Ts65Dn mice involved interactions between Erbb2 receptor tyrosine kinase 4 (*Erbb4*) with neuregulin1 (*Nrg1*), and *Erbb4* with neuregulin3 (*Nrg3*). We uncovered increased *Nrg1* to *Erbb4* signaling in OPCs and InNs, and decreased *Nrg3* to *Erbb4* signaling between InNs and OPCs in the Ts65Dn brain (Figures 3F and 3G). *Nrg1* regulates neuronal plasticity and migration, OL-lineage maturation, myelination, and dendritic arborization^88^. In InNs, *Nrg1* signaling through *Erbb4* plays a critical role in circuit development, neuronal differentiation and GABAergic transmission^89, 90^. Similarly, this signaling is vital in regulating OPC growth, proliferation, and differentiation into mOLs^91^. The dysregulation of this LR interaction between InNs and OPCs may thus contribute to DS cognitive disability attributable to excitatory/inhibitory (E/I) imbalance, modified neural network excitability, and reduced axon myelination. Like *Nrg1*, *Nrg3* is part of the neuregulin (*Nrg*) family and plays an important role in neural circuitry development through *Erbb4* signaling^92, 93^. Its downregulation has been implicated in several neurological and psychiatric conditions such as schizophrenia^94^. Importantly, the imbalance of Nrg signaling in the brain can drive neuropathology by compromising overall neural connectivity and circuit homeostasis: thus, the disrupted reciprocal interaction we observe in the Ts65Dn *Nrg1/3* LR networks may promote broad disruptions in brain architecture.

### Trisomy induces unique chromatin accessibility patterns in the Ts65Dncortex

To identify cell type-specific changes in the chromatin landscape of the Ts65Dn cortex, we analyzed differentially accessible regions (DARs) in snATAC-seq (Methods). We used DARs that met a FDR value of 5% to define statistically significant changes in chromatin-level expression between Ts65Dn and CTL offspring. We found that the majority (75.1%) of DARs were downregulated (Figure 4A) and were highly heterogeneous across cortical cell types (Figure 4B). To assess the genomic distribution of DARs, we used ChIPseeker^95^ (Methods). The distribution of DARs was relatively consistent across cell types, with a majority occurring in distal regulatory sites (Figure 4A). Across DARs, ∼6% were found in exons, ∼36% were found in promoters (<0-3kb), and ∼56% were found in distal regulatory sites (5’/3’ UTR, introns, >300kb downstream, or intergenic) (Figure 4C).

**Figure 4:**
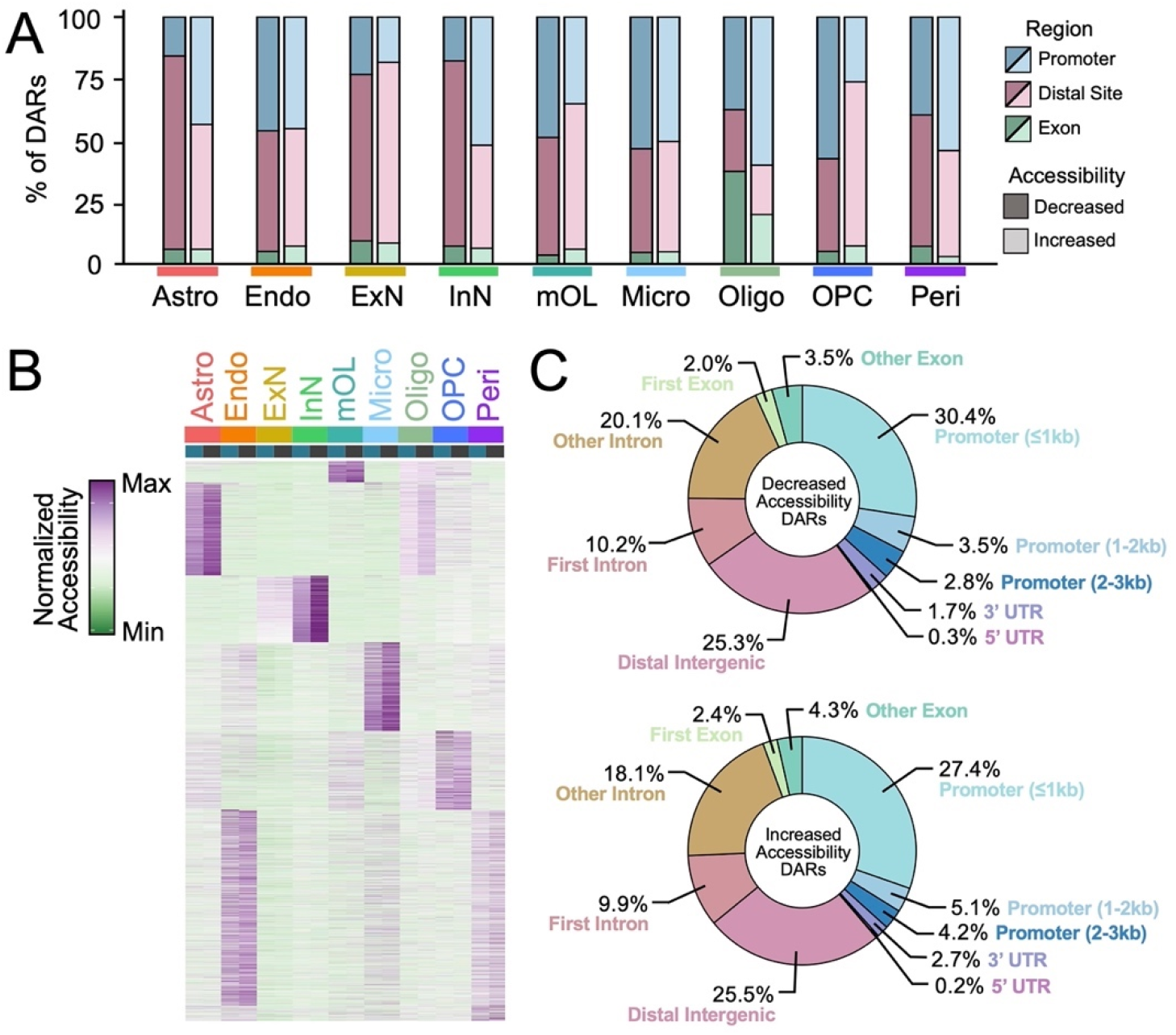
Distribution of cell type-specific chromatin accessible regions. (A) tacked barplot displaying the genomic distribution of differentially accessible regions (DARs). DARs located within promoters (< 0-3kb of the gene) are colored in blue, within distal regulatory sites (5’/3’ UTRs, introns, > 300kb downstream of the gene or intergenic) are colored in pink, or within exons are colored in green. Bars shaded in darker colors correspond to DARs showing decreased accessibility in Ts65Dn, and bars shaded in lighted colors correspond to DARs showing increased accessibility in Ts65Dn. Bars are grouped according to cell type identity. (B) Heatmap of the average number of cut sites within a DAR for each cell type by condition. Each column represents a cell type from Ts65Dn or CTL. The color code represents the row-normalized accessibility for each gene. (C) Pie chart of the genomic distribution of DARs that exhibit decreased or increased accessibility in Ts65Dn. Each fraction of the pie corresponds to a different genomic region, and is labeled according to the percentage of DARs associated with the specific genomic region.

Ts65Dn cells exhibited increased accessibility of ∼25% of Ts65Dn-encoded genes at the chromatin level. Similar to snRNA-seq, we observed heterogeneous accessibility levels of Ts65Dn triplicated genes across cell types, with only a few gene loci including the antioxidant *Sod1*, the kinase *Dyrk1a*, and the spliceosome component *Son* at significantly higher levels in at least 7 out of 9 identified cortical cell types (Figure 5A). Increased chromatin accessibility of *Olig1* and *Olig2* loci was observed selectively in astrocytes, mOLs, and OPCs. Overexpression of *Olig2* has been shown to preferentially drive progenitors towards Gfap+ reactive astrocytes rather than mOLs^96^ and can trigger transcriptional repression during myelinogenesis^97^. The *Runx1* locus exhibited increased accessibility in DS microglia; *Runx1* is a TF that modulates microglial gene expression during early postnatal life, but can increase in adults following brain injury or infection to promote microglial activation^98, 99^. The *Synj1* locus showed selective increased accessibility in DS InNs and is known for its involvement with endocytosis and synaptic vesicle cycling^100^. Specifically, elevated levels of *Synj1* have been shown to trigger deficits in age-dependent long-term memory retention in individuals with DS-related AD^101^.

**Figure 5:**
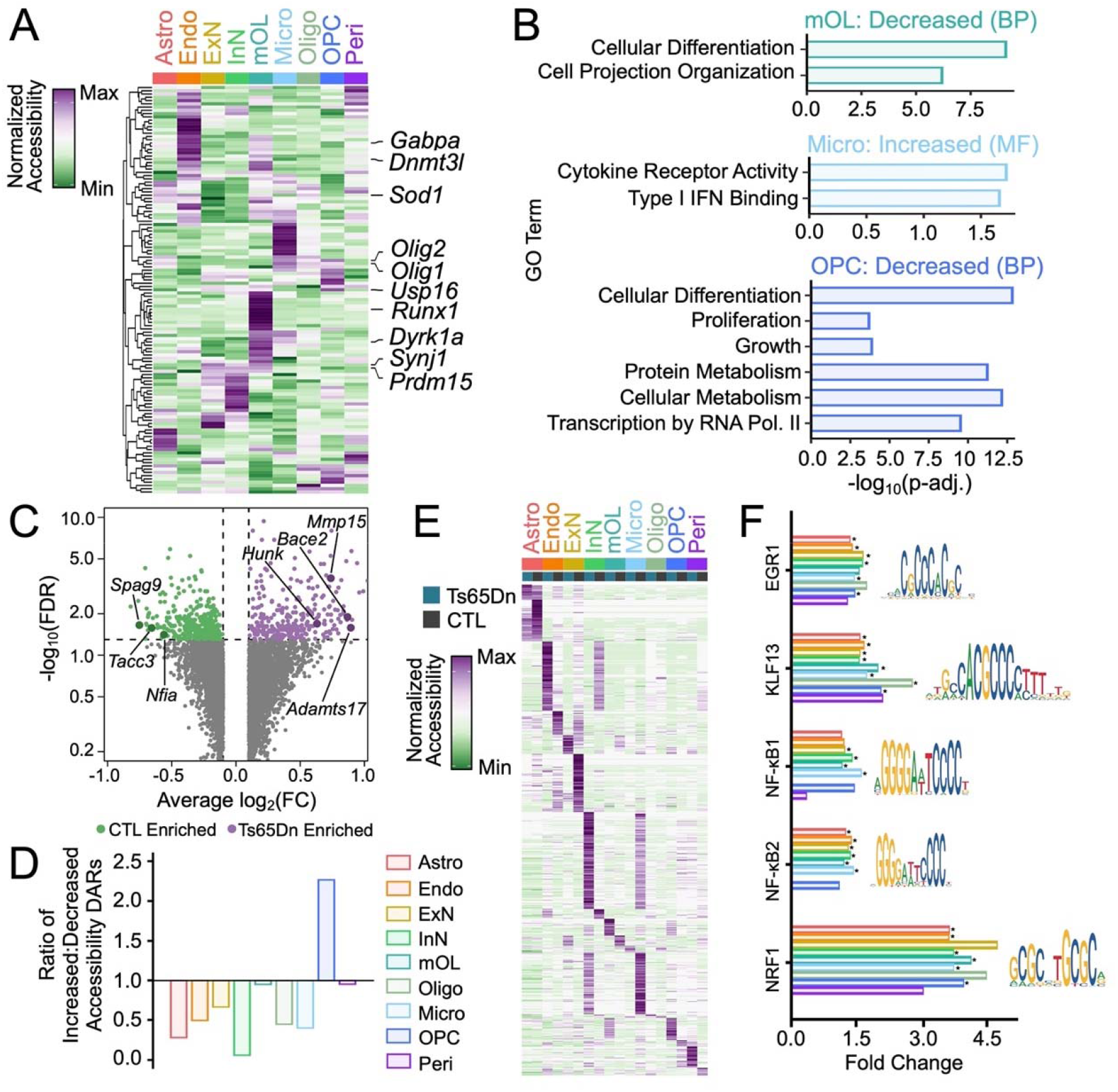
Chromatin accessibility landscape in the Ts65Dn brain (A) Hierarchically clustered heatmap of normalized gene accessibility for genes contained within the Ts65Dn chromosomal product. The plot displays gene activity across all cell types in snATAC-seq. Several genes of interest are indicated. Each column represents a cell type. The color code represents the row-normalized accessibility for each gene. (B) -log_10_(p-adj) value of enrichment for select gene ontology (GO) biological process (BP) or molecular function (MF) terms for mature oligodendrocytes (mOL), microglia (Micro) and oligodendrocytes precursor cells (OPC). Bars are colored according to cell type. (C) Volcano plot of DARs in Ts65Dn versus CTL conditions. Only genes with an average log_2_fold change (log_2_(FC)) > 0.1 or < -0.1. are included in the plot. Each dot represents a gene. Dots are colored according to enrichment: green dots are enriched in CTL, purple dots are enriched in Ts65Dn, and grey dots are not significantly enriched in either condition. The horizontal line depicts an FDR < 5%, such that all genes above this line are statistically significant. (D) Barplot displaying the ratio of DARs with increased to decreased accessibility across all cell types. A ratio of 1 indicates an equal number of increased and decreased DARs, while a ratio of greater or less than 1 indicates a greater number of increased or decreased DARs, respectively. Each bar is colored by cell type identity. Only statistically significant (FDR < 5%) are included. (E) Heatmap of average ChromVAR transcription factor (TF) motif activity for each cell type. Each column represents a cell type from Ts65Dn or CTL. The color code represents the row-normalized accessibility for each TF. (F) Barplot showing the relative fold change for select TFs between Ts65Dn and CTL. Each bar corresponds to a cell type. Bars labeled with asterisks indicate statistically significant enrichment (FDR < 5%) for the given TF. To the right of each grouped barplot is the position weight matrix (PWM) for the binding motif associated with each TF.

We next employed gene ontology (GO) analysis to assess biological pathways associated with chromatin accessibility changes in Ts65Dn mice (Methods)^102^. Among downregulated DARs from trisomic OPCs and mOLs, we observed an enrichment of GO categories associated with cell differentiation, projection organization, and regulation of metabolic processes (Figure 5B). We also found enrichment of biological process (BP) ontologies associated with growth and proliferation in trisomic OPC DARs with decreased accessibility, further suggesting a disruption in OPC development and a growth arrest typical of a senescence phenotype (Figure 5B). Consistent with the hyperactive inflammatory milieu in DS^14, 15^, we observed an enrichment of cytokine receptor activity and type I IFN binding amongst the top five enriched molecular function (MF) categories for increased accessibility DARs in Ts65Dn microglia (Figure 5B). Taken together, these findings demonstrate that trisomy has a profound genome-wide impact on gene expression, with a notable effect on genes related to inflammation and differentiation in glial cell populations.

To assess locus-specific changes in Ts65Dn OPC chromatin accessibility, we more closely examined population-level DARs. Herein, we observed reduced accessibility at several cell growth and mitosis regulatory loci (Figure 5C), including *Tacc3*, *Knstrn*, *Ptprg*, and *Spag9*, whose suppression can induce cell cycle arrest and tend cells towards a senescent state^103–106^. Further, we observed increased accessibility at several CSPG loci, including *Tnr*, *Bcan*, *Vcan*, *Acan*, and *Ncan*^107^ (Figure 5C). CSPGs have been found to negatively regulate OPC proliferation, differentiation and remyelination, as well as induce production of pro-inflammatory cytokines^108, 109^. Similar to our findings in snRNAseq data, we found that Ts65Dn OPCs exhibited an increase in accessibility at several endoprotease loci, including *Mmp15*/*19*, *Adam12*, and *Adamts5*/*17* loci (Figure 5C), all of which play a role in modulating neuroimmune activity through the regulation of pro-inflammatory cytokine availability and signaling^73, 74^.

Structural changes within and around senescent cells are also accompanied by dramatic molecular changes to the epigenetic landscape, including a global reduction in heterochromatin^110–112^. To investigate whether cortical Ts65Dn OPCs exhibited this senescence-associated loss of heterochromatin, we compared the number of DARs with increased or decreased accessibility in Ts65Dn versus euploid CTL across all cell types. We found a greater number of peaks with increased accessibility in Ts65Dn relative to CTL only in OPCs, consistent with a net increase in chromatin accessibility (Figure 5D). All other cell types profiled through snATAC-seq exhibited a greater number of peaks with decreased accessibility, suggestive of a cell type-specific loss of heterochromatin consistent with selective senescence in Ts65Dn OPCs.

### Cell type-specific transcription factors in the trisomic mouse cortex

To gain insight into transcription factor (TF)-mediated gene regulation in Ts65Dn mice, we constructed cell type-specific TF regulatory networks. We used chromVAR^113^ to identify shared and cell type-specific gene-chromatin interactions based on snATAC-seq and observed unique TFs enriched across all cortical cell types (Figure 5E; Methods).

ChromVAR detected enrichment of the early growth response factor 1 (EGR1) TF binding site in DS InNs, which was supported by increased *Egr1* transcription in Ts65Dn InNs. EGR1 regulates the development of GABA receptor subunit genes, thus acting as a key regulator of InN development^114^ (Figure 5F). A similar pattern was observed for KLF13, which binds to regulatory regions of genes that are important for OL differentiation and maturation, such as *Mag*, *Mbp*, and *Plp1*^115^. There was increased accessibility at the KLF13 binding motif in Ts65Dn mOLs, as well as increased transcript level expression of *Klf13* detected by snRNA-seq (Figure 5F).

In trisomic endothelial cells, ExNs, InNs, microglia, and mOLs, we observed a marked increase in motif enrichment for nuclear factor kappa-light-chain-enhancer of activated B cells (NF-κB1 and NF-κB2), a critical modulator of cell growth, cell survival, development, and inflammation^116^ (Figure 5F). This increase in NF-κB1/2 TF accessibility across several cortical cell types suggests an increase in inflammatory signaling, consistent with the hyperimmune milieu observed in DS. Additionally, we found an enrichment of nuclear respiratory factor 1 (NRF1) motif accessibility in all trisomic major neuronal and non-neuronal cell types except for mOLs and pericytes (Figure 5F). NRF1 is a key regulator of cell growth, apoptosis, senescence, neurogenesis, genomic instability, and mitochondrial function^117^. A disruption in mitochondrial biogenesis, driven by NRF1 dysregulation, has been observed in many neurodegenerative and neurodevelopmental disorders including AD, Parkinson’s Disease (PD), and Huntington’s Disease (HD), and may likewise play a role in widespread neurogenesis or myelination deficits in DS^117–119^. Lastly, we noted an enrichment of the hairy and enhancer of split-1 (HES1) motif in Ts65Dn ExN, InN, and OPCs relative to euploid CTL (Figure 5F). HES1 represses proneural or differentiation genes to keep stem cell progenitors in quiescent or proliferative states^120–122^. Importantly, HES1 expression is downregulated during myelination, and HES1 overexpression in OPCs results in impaired oligodendrogenesis^123^. Together, these results show that Ts65Dn trisomy leads to widespread perturbation of TF activity and binding motif accessibility, which may contribute to key elements of trisomy-associated neuropathology.

### Deep layer cortical Ts65Dn OPCs display increased senescence

To support our finding of an enriched senescence-associated gene signature in Ts65Dn OPCs, we assayed coronal brain sections from 3mo and 6mo mice for lysosomal senescence-associated-β- gal (SA-β-gal) activity coupled with immunohistochemistry (IHC; Methods). We profiled several major cortical cell types using antibodies against canonical marker genes, including microglia (Iba1+), astrocytes (Aqp4+), OL-lineage cells (Olig2+), OPCs (Pdgfrα+), and neurons (NeuN+). Ts65Dn mice exhibited a 1.7- and 1.8-fold increase in Olig2+ SA-β-gal activity at 3mo and 6mo, respectively, as well as a 1.4- and 1.3-fold increase in Pdgfrα+ SA-β-gal activity at 3mo and 6mo, respectively (Figures 6A and 6B). In contrast, astrocytes, neurons, and microglia did not demonstrate a significant increase in SA-β-gal activity in Ts65Dn mice (Figure 6C). Intriguingly, we observed that the increase in OL-lineage senescence was restricted to the deep layers (L5-6) of the Ts65Dn cortex and was absent in the upper cortical layers (L1-4) and the corpus callosum (CC) (Figure 6D), suggestive of regional specificity.

**Figure 6:**
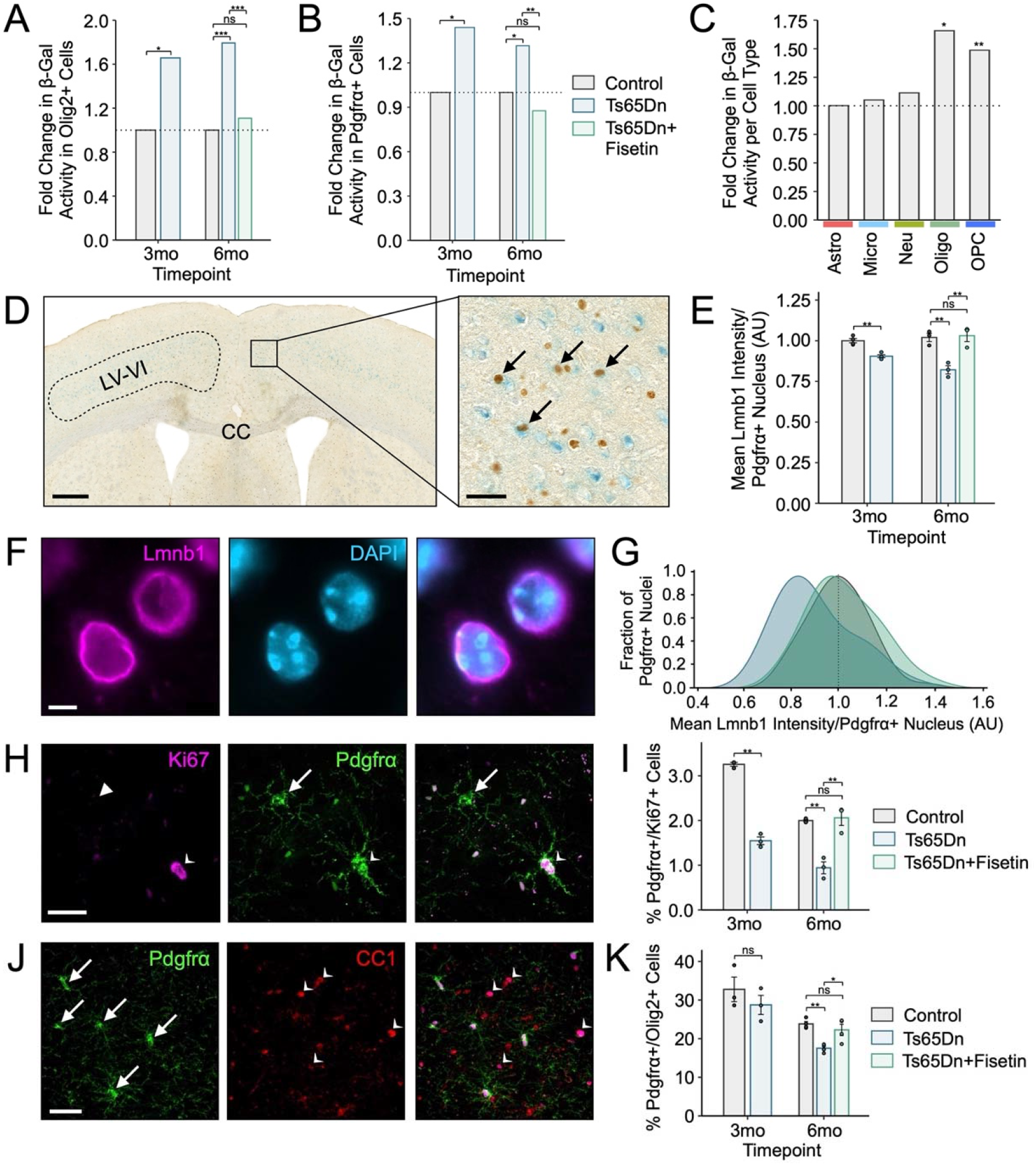
Deep layer cortical Ts65Dn OPCs exhibit hallmarks of senescence, which are rescued by Fisetin treatment. (A) Barplot of SA-β-gal activity in Olig2+ cells (OL-lineage) in 3mo and 6mo mice, showing a significant increase in SA-β-gal+/Olig2+ cells in Ts65Dn at both timepoints, with a rescue in 6mo Ts65Dn mice treated with Fisetin. Bars represent the average values for each condition (*n* = 3 mice per condition, *n* = 3 replicates per mouse), and dots represent the average values for each mouse per condition. The *y*- axis represents the SA-β-gal intensity in Ts65Dn or Ts65Dn+Fisetin mice to CTL mice; the horizontal dashed line intercepts the *y*-axis at 1, indicating normalized SA-β-gal activity to CTL at both the 3mo and 6mo timepoints. Significance is determined using the two-tailed Student’s t-test at 3mo, or using the ANOVA test at 6mo. *p-value < 0.05; **p-value < 0.01, ***p-value < 0.001, ns: not significant; error bars represent the average ± 1 standard deviation (SD). (B) as in (A) but depicting SA-β-gal activity in Pdgfrα+ cells (OPCs), showing a significant increase in SA-β-gal+/Pdgfrα+ cells in Ts65Dn at both timepoints, with a rescue in 6mo Ts65Dn mice treated with Fisetin. (C) Barplot of the fold change in SA-β-gal activity across several major cortical cell types in 3mo Ts65Dn versus CTL mice. Astro, astrocytes; Micro, microglia; Neu, neurons; Oligo, oligodendrocytes; OPC, oligodendrocyte precursor cells. Only Oligos and OPCs exhibit statistically significant increases in SA-β-gal activity in Ts65Dn versus CTL, as depicted through an asterisks above the bar. (D) SA-β-gal staining in coronal brain sections from a 3mo Ts65Dn mouse; arrows point to Olig2+ (brown) SA-β-gal+ (blue) cells throughout the cortex. Panel on the right shows a high magnification image of the area demarcated in the left panel. Images are representative of those observed in samples from Ts65Dn and CTL mice (*n* = 3 mice per condition, *n* = 3 replicates per mouse). Scale bars are shown in the bottom left corner of each panel: left panel, 500μm; right panel, 50μm. (E) Barplot of LaminB1 (Lmnb1) intensity in arbitrary units (AU) per OPC (Pdgfrα+) nucleus. Values have been normalized to the mean intensity found in CTL replicates. Bars represent the average values for each condition (*n* = 4 mice per condition at 3mo and *n* = 3 mice per condition at 6mo, *n* = 3 replicates per mouse), and dots represent the average values for mouse per condition. (F) Immunostaining for Lmnb1 (pink) and nuclear DAPI (blue) in a 3mo Ts65Dn mouse. Images are representative of those observed in samples from Ts65Dn and CTL mice (*n* = 3 mice per condition, *n* = 3 replicates per mouse). Scale bar is shown in the bottom left corner of the panel: 5μm. (G) Density plots showing Lmnb1 intensity distribution per OPC (Pdgfrα+) nucleus in 6mo mice; bin populations were calculated by averaging the relative contribution per sample. (H) Immunostaining for Ki67 (pink) and Pdgfrα (green) in a 3mo Ts65Dn mouse. Images are representative of those observed in samples from Ts65Dn and CTL mice (*n* = 3 mice per condition, *n* = 3 replicates per mouse). Arrow points to Pdgfrα+ cells, and arrowheads point to Ki67+/Pdgfrα+ cells. Scale bar is shown in the bottom left corner of the panel: 20μm. (I) Barplot of the percentage of Ki67+/Pdgfrα+ cells (proliferating OPCs), showing a significant increase in proliferating OPCs cells in Ts65Dn at both timepoints, with a rescue in 6mo Ts65Dn mice treated with Fisetin. Bars represent the average values for each condition (*n* = 3 mice per condition, *n* = 3 replicates per mouse), and dots represent the average values for each mouse per condition. (J) Immunostaining for CC1 (red) and Pdgfrα (green) in a 3mo Ts65Dn mouse. Images are representative of those observed in samples from Ts65Dn and CTL mice (*n* = 3 mice per condition, *n* = 3 replicates per mouse). Arrow points to Pdgfrα+/Olig2+ cells, and arrowheads point to CC1+/Olig2+ cells. Scale bar is shown in the bottom left corner of the panel: 15μm. (I) Barplot of the percentage of Ki67+/Pdgfrα+ cells (proliferating OPCs), showing a significant increase in proliferating OPCs cells in Ts65Dn at both timepoints, with a rescue in 6mo Ts65Dn mice treated with Fisetin. Bars represent the average values for each condition (*n* = 3 mice per condition, *n* = 3 replicates per mouse), and dots represent the average values for each mouse per condition.

To confirm these findings, we analyzed 3mo and 6mo coronal sections for additional hallmarks of senescent cells: reduction of nuclear envelope protein LaminB1 (Lmnb1) intensity and reduction of proliferation^66, 124^. Lmnb1 decline associated with senescence can be detected at the protein-level through immunofluorescence (IF) staining and subsequent measurement of fluorescence intensity. We thus stained for Lmnb1 in Ts65Dn and euploid CTL tissues in a cell type-specific manner (using Pdgfrα to stain for OPCs) and compared Lmnb1 fluorescence intensity between conditions (Methods). As expected, we observed a decline in cortical OPC (Pdgfrα+) Lmnb1 intensity by 10.4% at 3mo and 14.4% at 6mo (Figure 6E–6G). Senescent cells are also typified by an irreversible cell cycle and proliferation arrest^66^: proliferating cell populations can be identified by quantifying nuclear Ki67 positivity (Methods). We observed a 51.7% and 58.0% reduction in proliferating Ki67+ OPCs in the Ts65Dn cortex at 3mo and 6mo, respectively (Figures 6H and 6I), suggestive of a growth arrest typical of senescence in DS OPCs. Importantly, OPCs localized to the CC did not exhibit a change in Ki67 positivity in DS (Figure 6–figure supplement 1A), thus supporting a selective regional and cell type specific senescence in cortical DS OPCs.

To determine how OPC senescence affects cortical populations of oligodendrocyte lineage cells, we quantified changes in OPC and mOL cell counts in coronal brain sections from Ts65Dn versus euploid littermate CTLs. Prior studies have shown that trisomy-associated triplication of Olig2 results in a transient early-life increase in OPCs, followed by a progressive decline in the progenitor pool with a resultant deficit in myelination that persists into adulthood^7, 96^. We stained for Olig2 to mark the OL-lineage, and Pdgfrα or CC1 to identify OPCs or mOLs, respectively. At both the 3mo and 6mo timepoint, we discovered a decline in the percentage of Olig2+/Pdgfrα+ cells in the cortex, although only that at the 6mo timepoint met statistical significance (Figures 6J and 6K). Despite changes in OPC proliferation and abundance in the trisomic cortex, the percentage of Olig2+/CC1+ mOLs was not significantly changed in the cortex of Ts65Dn mice (Figure 6 – figure supplement 1B). Likewise, we found no change in cortical Mbp mRNA levels through quantitative PCR (qPCR) (Figure 6–figure supplement 1C), or in Mbp fluorescence intensity measured through immunostaining (Figure 6–figure supplement 1D). Overall, this analysis suggests that OPC senescence alters progenitor abundance, but not OL maturation or myelination in the Ts65Dn cortex.

### Treatment with the senescence-reducing flavonoid Fisetin rescues cellular and behavioral deficits in Ts65Dn mice

Senescent cell anti-apoptotic pathways (SCAPs) are a key feature of senescent cells, allowing them to evade apoptotic cell death^66^. SCAPs can be targeted by small molecule senolytic compounds to selectively clear senescent cells from pathological tissues^125^. Fisetin, a natural flavonoid, has senolytic properties and can be added to the diet as a supplement with minimal adverse effects^126^. We fed Ts65Dn mice a diet supplemented with 500ppm Fisetin, as previously described^127^ (Methods). As expected, we observed a significant reduction in SA-β-gal positivity in both Pdgfrα+ and Olig2+ cells in the cortex of trisomic mice given Fisetin (Figures 6A and 6B). Fisetin supplementation also restored Lmnb1 protein levels in Ts65Dn OPCs to CTL levels (Figure 6E),increased the proportion of Ki67+/Pdgfrα+ cells in the Ts65Dn cortex (Figure 6I), and restored the abundance of cortical OPCs (Figure 6K). While the number of OPCs was observed to significantly decrease in the 3mo Ts65Dn CC, there was no significant difference in the number of OPCs in the CC at 6mo, and no effect of Fisetin treatment on CC OPC cell counts (Figure S2E). Likewise, there was a significant decrease in the number of Ki67+/Pdgfrα+ at both 3mo and 6mo, but no rescue of this reduction in proliferation activity through Fisetin treatment at 6mo in the CC (Figure S2F), consistent with a region-specific effect of senescent OPCs in the Ts65Dn cortex.

DS mouse models demonstrate increased microglial and astrocyte reactivity, as well as increased inflammatory cytokine levels and interferon signaling^14–17, 20, 60–62^. Given that senescent cells modify the local niche inflammatory environment through the SASP^76^, we hypothesized that Fisetin administration and resulting reduction in OPC senescence would alter local microglial phenotype. To begin, we counted the number of Iba1+ microglia in the cortex of 3mo and 6mo Ts65Dn and CTL mice. We observed a significant increase in the number of Iba1+ microglia in trisomic mice at both time points, and Fisetin administration was associated with a restoration of Iba1+ microglia at 6mo (Figure 7A). Next, we tested whether the activation state of microglia was altered in Ts65Dn mice by quantifying the abundance of the phagocytic marker Cd68 within Iba1+ cells. Interestingly, we observed a decrease in the proportion of Cd68+/Iba1+ cells in Ts65Dn at 6mo (Figures 7B and 7C), suggestive of a reduction in phagocytic microglial activity. This decline in microglial phagocytic capacity is consistent with prior reports demonstrating that aged microglia display a reduction or dysfunction in phagocytosis^128–130^, thus suggesting that the Ts65Dn cortex and its associated senescent microenvironment may contribute to an impairment in microglia phagocytic activity, rescuable by Fisetin treatment.

**Figure 7:**
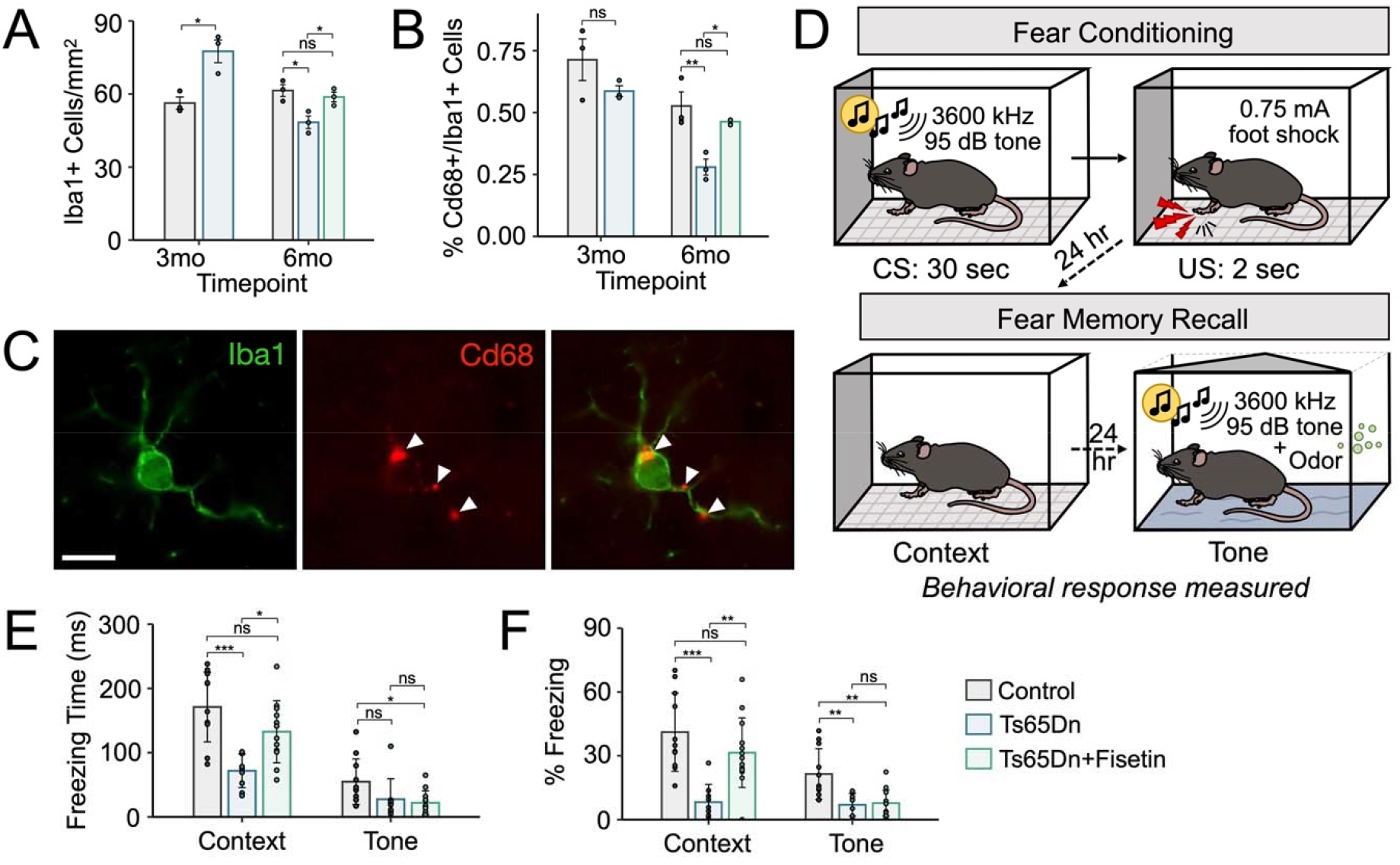
Fisetin rescues microglial phenotype and fear memory behavior in Ts65Dn mice. (A) Barplot of the number of Iba1+ (microglia) cells per mm^2^ in the cortex in 3mo and 6mo mice, showing a significant increase in microglia in Ts65Dn at both timepoints, with a rescue in 6mo Ts65Dn mice treated with Fisetin. Bars represent the average values for each condition (*n* = 4 mice per condition, *n* = 3 replicates per mouse), and dots represent the average values for each mouse per condition. Significance is determined using the two-tailed Student’s t-test at 3mo, or using the ANOVA test at 6mo. *p-value < 0.05; **p-value < 0.01, ***p-value < 0.001, ns: not significant; error bars represent the average ± 1 standard deviation (SD). (B) as in (A), but depicting the percentage of Cd68+/Iba1+ (activated microglia) in the cortex, showing a significant decrease activated microglia at 6mo, with a rescue in 6mo Ts65Dn mice treated with Fisetin. (C) Immunostaining for Iba1 (green) and Cd68 (red) in a 3mo Ts65Dn mouse. Images are representative of those observed in samples from Ts65Dn and CTL mice (*n* = 3 mice per condition, *n* = 3 replicates per mouse). Arrows point to Cd68+ puncta in Iba1+ microglia. Scale bar is shown in the bottom left corner of the panel: 15μm. (D) Schematic representation of fear conditioning and memory recall behavioral battery. *n* = 9 biological replicates per condition. (E) as in (A), but depicting the time to freezing behavior depending on context and tone, showing a significant decrease in context-based but not tone-based freezing in Ts65Dn mice, with a rescue in the context-based behavior in 6mo Ts65Dn mice treated with Fisetin. Bars represent the average values for each condition (*n* = 9 mice per condition), and dots represent the data collected for each mouse per condition. (F) as in (E), but depicting the percentage of context-related freezing behavior, showing a significant decrease in context and tone-based freezing in Ts65Dn mice, with a rescue in the context-, but not tone-based behavior in 6mo Ts65Dn mice treated with Fisetin.

Individuals with DS, as well as DS mouse models, exhibit pronounced learning and memory deficits that worsen with age. We thus wanted to determine whether senescence-reducing Fisetin supplementation treatment could rescue learning-associated behavioral deficits in 6mo DS mice. To do so, we employed a contextual fear conditioning paradigm, as previously done for Ts65Dn mice^49^, which pairs a foot shock (unconditioned stimulus, US) with a context (conditioned stimulus, CS), and subsequently tests memory strength by measuring fear-associated freezing behavior in 6mo mice exposed to the CS after 24-hours (Figure 7D; Methods). We found, as expected, that DS mice exhibited significantly lower freezing behavior compared to CTL littermates in both contextual and tone associated freezing behavior (Figure 7E and 7F). Contextual, but not tone-associated freezing behavior, was rescued in 6mo DS mice treated with Fisetin, indicating improved associative learning (Figure 7E and 7F). Overall, the behavioral data appears to suggest that, in Ts65Dn mice, cognitive impairment may be caused, at least in part, by cortical OPC senescence, and that this behavioral impairment is rescuable by Fisetin treatment.

## Discussion

The present study leverages an integrated transcriptome and chromatin accessibility landscape of the Ts65Dn mouse cortex to identify trisomic OPCs that undergo accelerated senescence. The study characterizes cell type-specific effects of trisomy and dosage compensation, identifying putative mechanisms of cell-cell signaling dysfunction in the mature brain. Importantly, we assemble multiple lines of evidence supporting OPC senescence in Ts65Dn mice, and we find that OPC senescence is spatially concentrated in deeper cortical layers. This suggests a unique phenotype for cortical gray matter OPCs and a role for the cortical niche in programming OPC state, similar to what has been recently reported for other non-neuronal cells^131^.

Our work builds upon previous data suggesting cell-autonomous defects in OL maturation in mouse models of DS^7^, as our data confirms a reduction in OPC proliferation and fewer mOLs in the CC. In the developing DS brain, there is a cell fate shift from neurogenesis to gliogenesis, resulting in a transient increase in Olig2+ cells, followed by a decrease in mOLs and an increase in astrocytes with age, which is thought, in part, to be driven by triplication of OLIG1/2 TFs^132^. This supports a model of defective OL-lineage differentiation capacity and a shift in glial cell commitment. Importantly, our data show that at 6mo, Ts65Dn mice show a reduction in cortical gray matter OPCs, but not CC OPCs, suggesting an age-restricted cortical gray matter phenotype. Therefore, DS OPCs are uniquely vulnerable to impaired maturation, but the local cortical microenvironment may be important as a second hit to promote senescence.

There is growing recognition that OPCs are an important mediator of neuroinflammation and aging phenotypes in the brain. OPCs exhibit regional, transcriptional, and functional heterogeneity, including differences in calcium signaling and electrical activity^133, 134^. OPCs become more regionally diverse, acquire ion channels^130^, and undergo an inflammatory transformation with age^135^. OPCs harbor a highly dynamic cytokine receptor-mediated surveillance mechanism that responds to cues of injury and inflammation^136^, and these receptors promote neuroinflammatory responses upon IFN signaling. OPC signaling to the microenvironment is bidirectional: local inflammation influences OPC ability to proliferate and differentiate, while OPCs also exert an immunomodulatory function on neighboring cells. Therefore, our study adds to this existing literature to suggest a region-specific disruption in OPC function with aging in DS.

The DS brain exhibits several pathologic changes, including DNA damage, chromatin instability, oxidative stress, mitochondrial dysfunction, and immune activation, that may predispose to cellular senescence. In particular, there is a chronic hyper-interferon state in the DS brain due to the increased gene dosage of IFNARs^16, 137–139^. IFNAR inhibition reverses senescent microglia phenotype in human DS cells, and chronic interferon-opathy may contribute to accelerated aging and neurodegeneration in DS^139^. Nevertheless, the precise gene-environment mechanisms that regulate region-specific OPC senescence, at least in Ts65Dn mice, are unclear. Interestingly, our results are consistent with previous studies of the aged AD-affected brain, in which inflammatory Aβ plaque-associated OPCs were found to exhibit increased senescence^140^. This suggests a link between OPC senescence and a neuroinflammatory microenvironment in aging. Other investigators have reported senescence in human DS neural progenitor cells (NPCs) and microglia, but these studies were done in human cell culture models^141^. Our findings are exclusively derived from the Ts65Dn mouse model, and this does not exclude neural progenitor or microglia senescence at other developmental or stimulus-dependent contexts. Further validation of these findings in human tissue across the lifespan will be critical to understanding the role of senescence in individuals with DS.

There are several limitations to the present study. Droplet-based single cell genomics is limited by gene and chromatin fragment “drop-out” and is sparser than bulk approaches. Single-cell RNA-seq is biased against lowly expressed genes, which includes many immune molecules and senescence associated transcripts in the brain. Chromatin accessibility may be a coarse indicator of senescence associated epigenetic changes in OPCs, and a detailed analysis of histone landscape and 3D genome architecture of DS OPCs is warranted. Furthermore, gene expression levels may not correlate with protein levels, particularly with aging, and our data do not reflect possible post-transcriptional regulatory mechanisms in translational control. Ts65Dn mice do not exhibit amyloid plaques consistent with human AD^142^. The use of the Ts65Dn mouse model also harbors inherent limitations, namely the inclusion of only ∼55% of HSA21 protein-coding orthologs^143^. Indeed, Ts65Dn mice do not triplicate many of the trisomic human loci, while also creating trisomic-imbalance for several MMU17 genes unrelated to HSA21. A recent study conducted on a newly-developed DS model – Ts66Yah – demonstrates that several genes located on MMU17 that are triplicated in Ts65Dn mice may play a role in DS-associated neuropathology and behavior^144^. Conclusions drawn from use of the Ts65Dn mouse model of DS must therefore be interpreted and understood in the context of existing limitations of the model, and warrant supplementation in other murine contexts or in human DS tissue for increased validity and translatability. Further, senescence-reducing therapy with Fisetin is not specifically targeted to OPCs, and the effects of Fisetin may be related to non-cell-autonomous mechanisms. This does not exclude the possibility of direct effect of Fisetin on other cell types. Future work using transgenic models to selectively deplete senescent OPCs will be important to demonstrate the importance of this phenomenon to DS-associated cognitive decline and neurodegeneration.

Despite these limitations, our study identifies a cell type-specific senescence phenotype in Ts65Dn OPCs that is targetable with senescence-reducing therapies. These findings shed light on potential drivers of chronic inflammation and cognitive decline in the DS brain. Furthermore, the integrated single-nucleus transcriptomic and chromatin landscape in the Ts65Dn brain is an important resource for discovery in DS neurobiology.

## Materials and Methods

### Key Resources Table

**TABLE 1.**
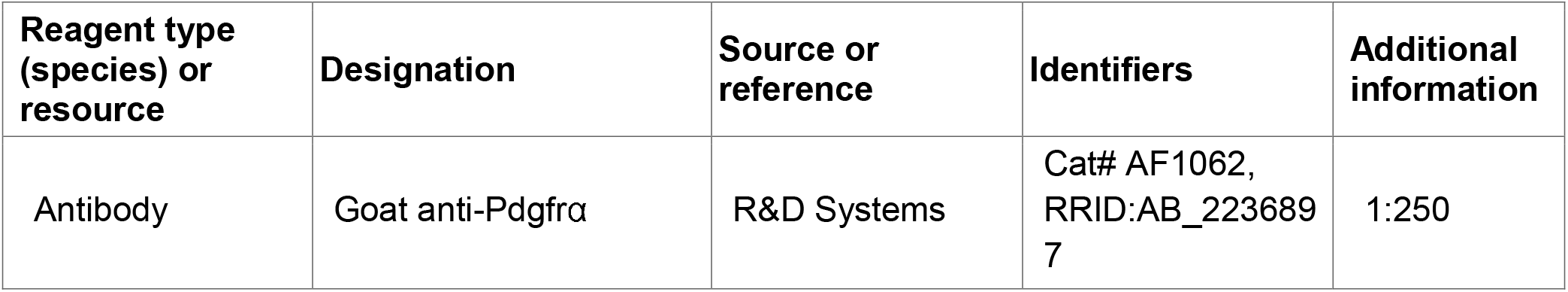
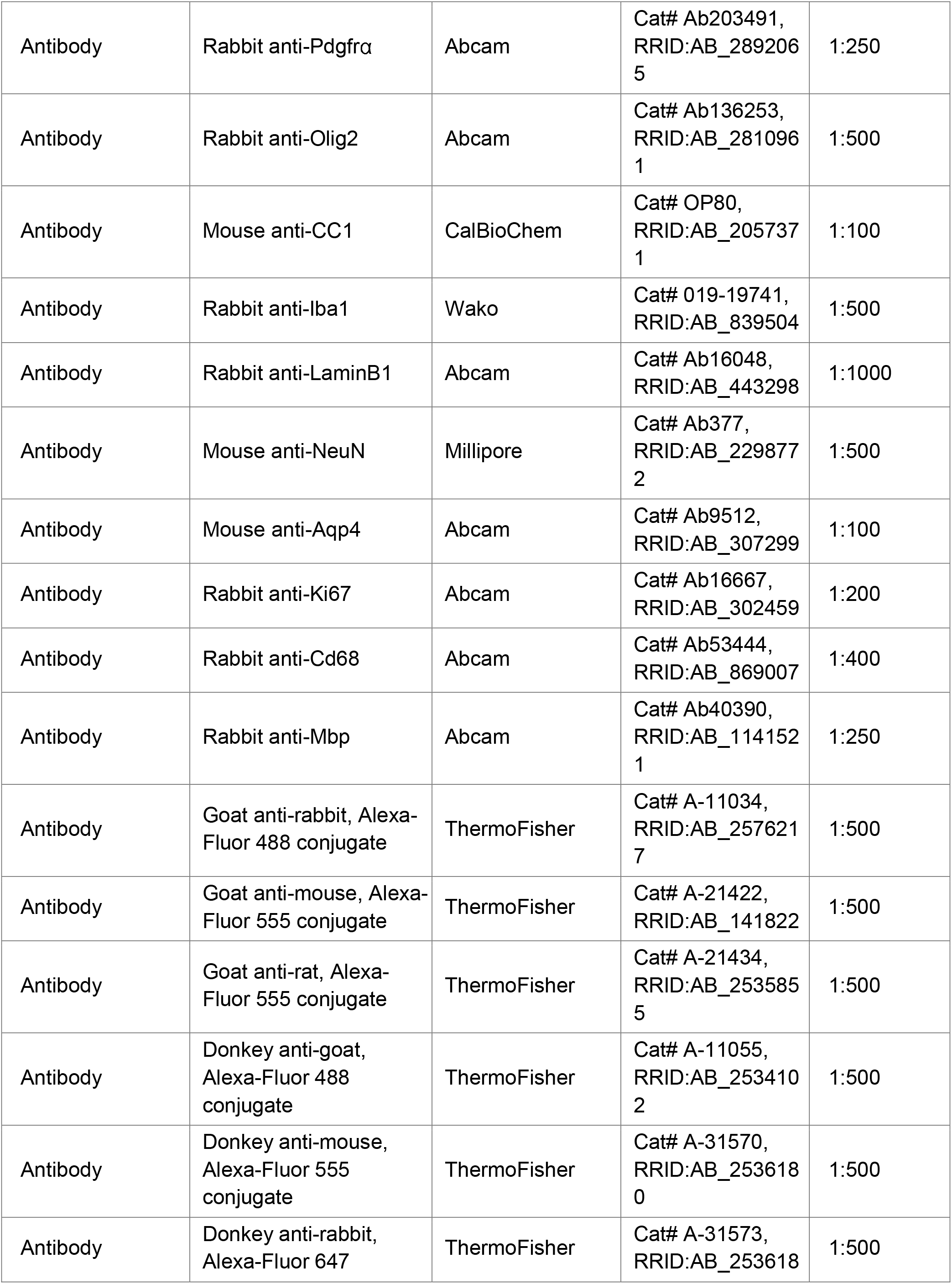
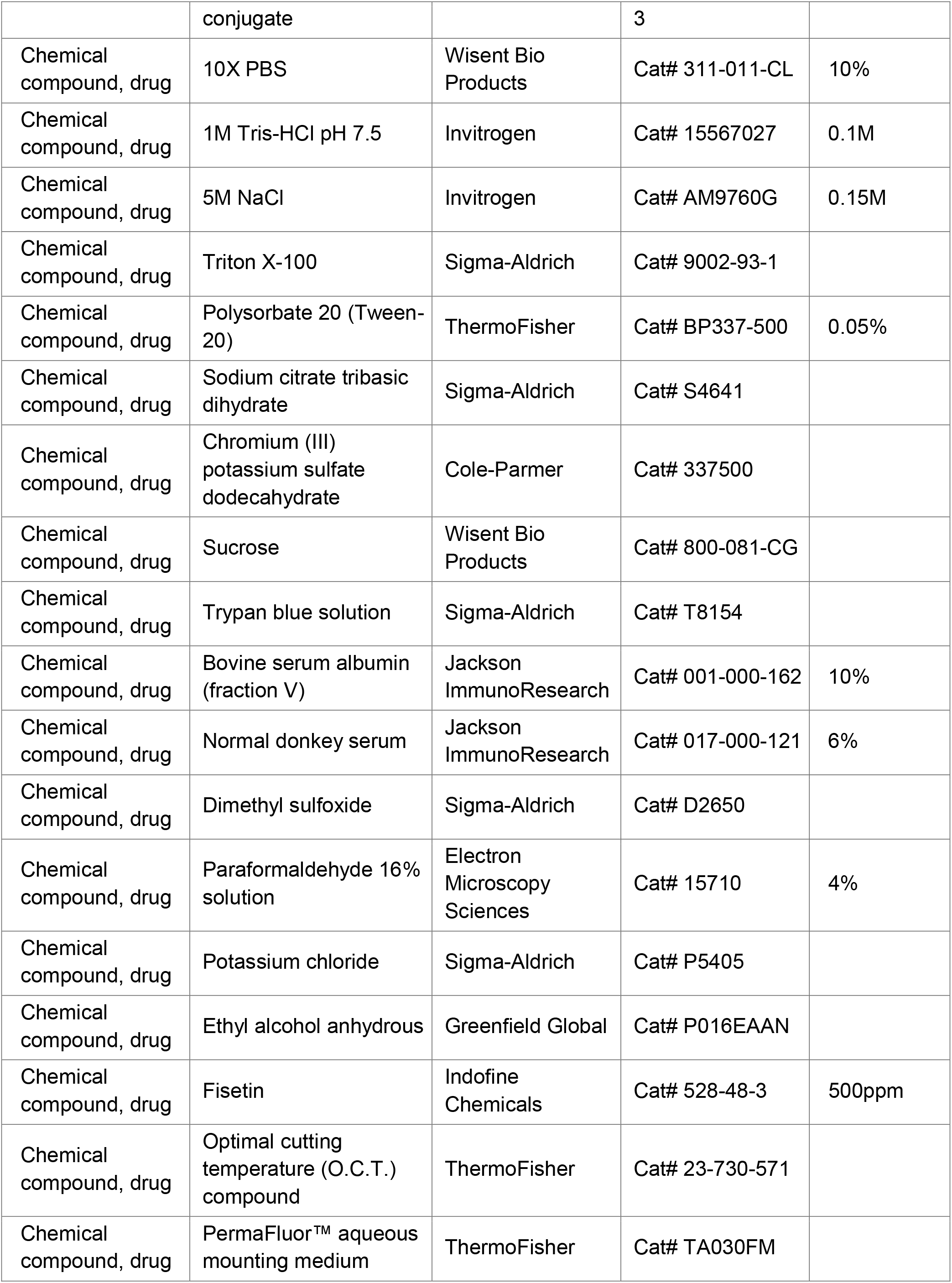
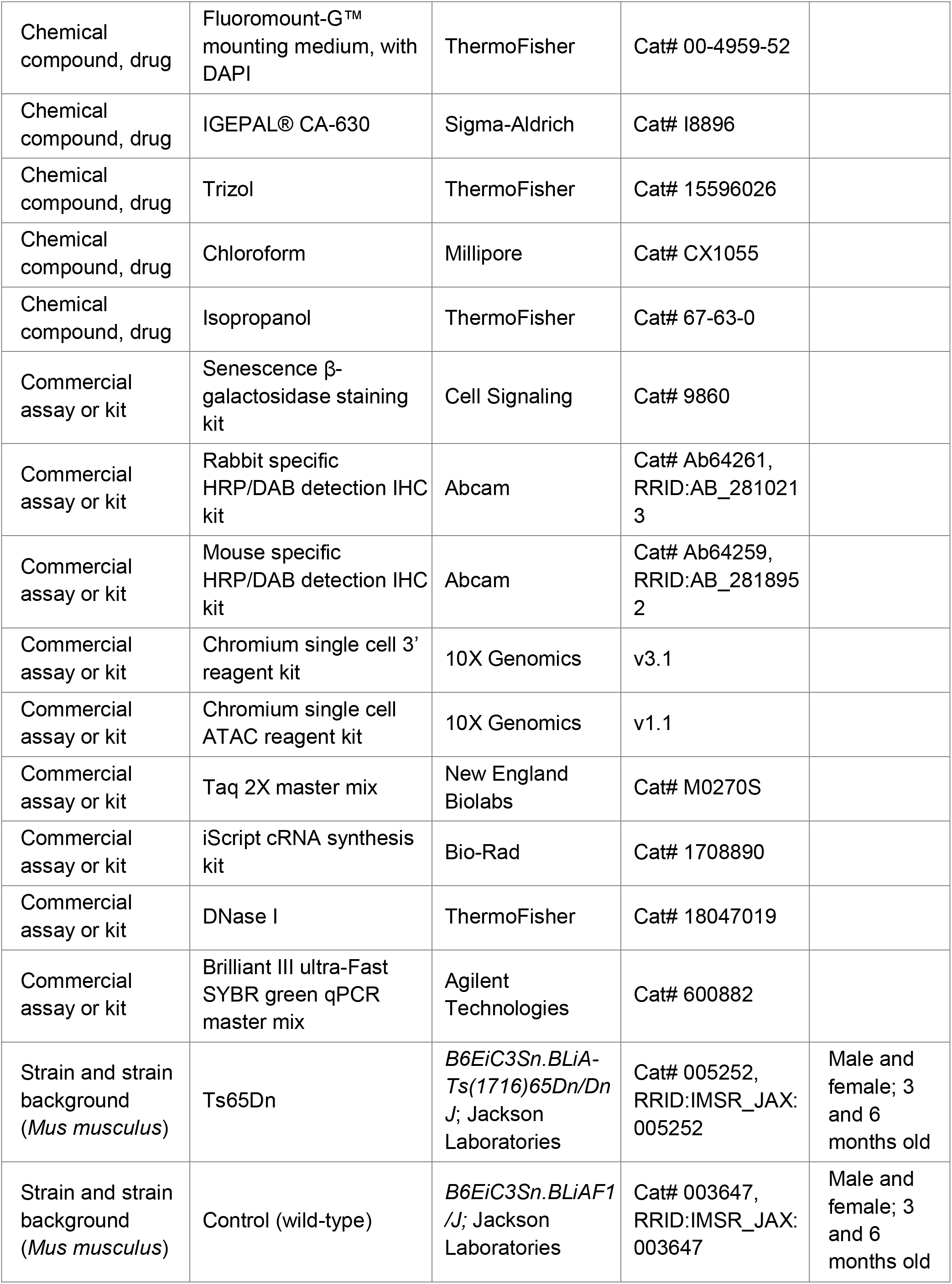
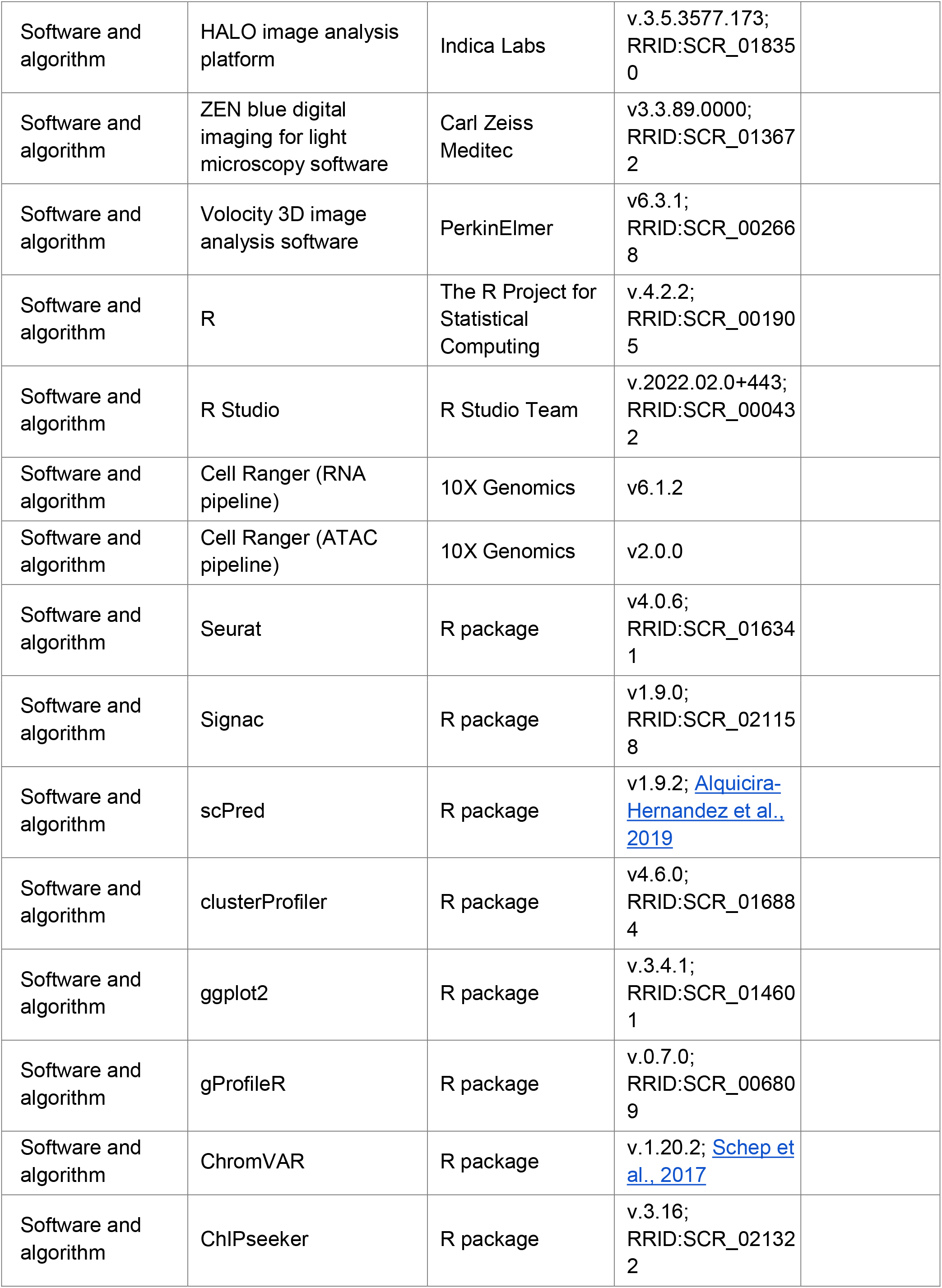
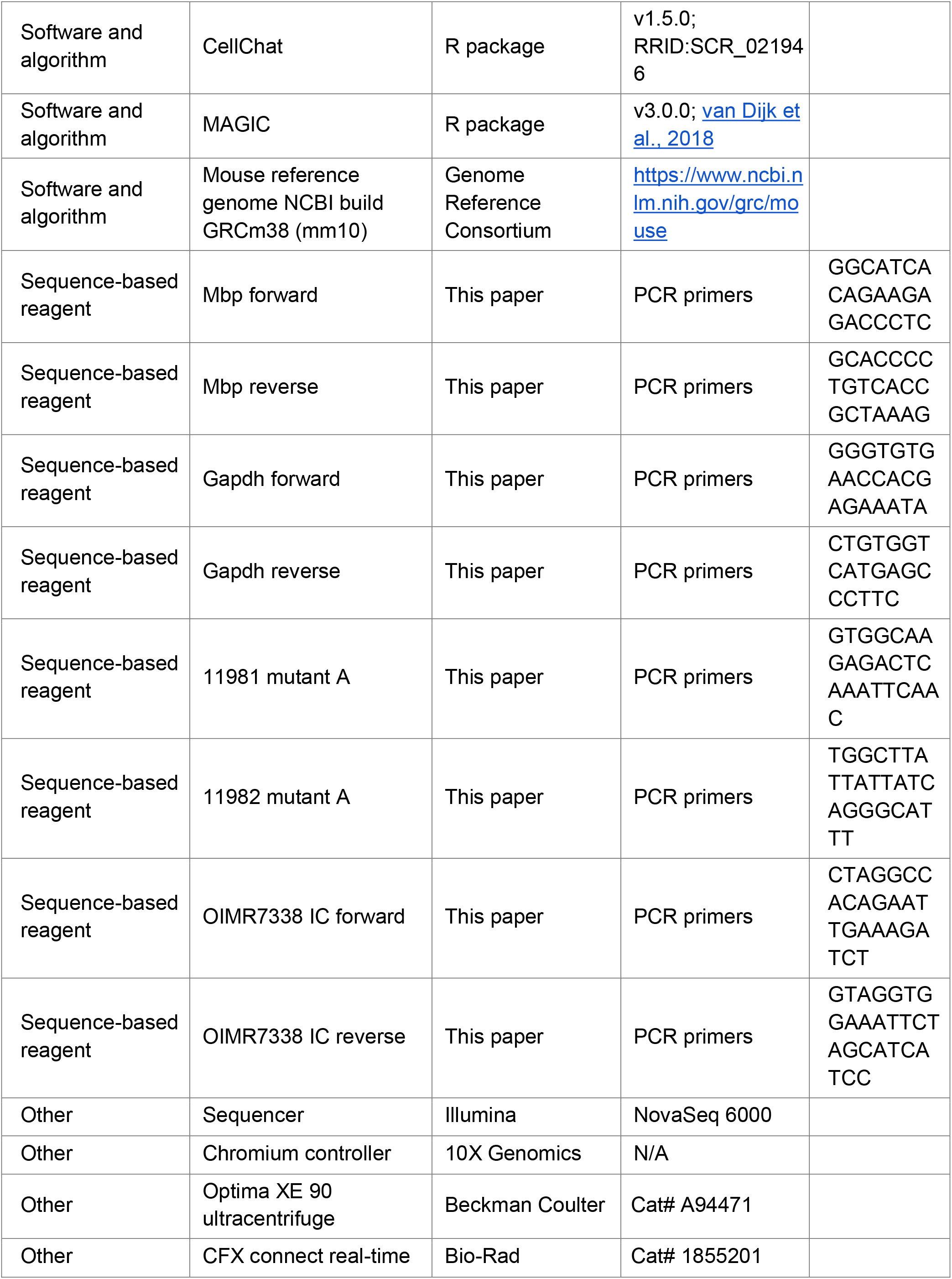
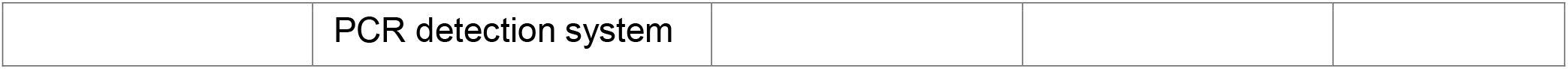

### Mice

All animal experiments were performed in accordance with the Canadian Council of Animal Care policies. Ts65Dn (*B6EiC3Sn.BLiA-Ts(1716)65Dn/DnJ*, RRID:IMSR_JAX:005252) and euploid controls (*B6EiC3Sn.BLiAF1/J*, RRID:IMSR_JAX:003647) were purchased from Jackson Laboratories. Ts65Dn mice were bred with euploid controls as recommended by Jackson Laboratories. The animals were kept in 12 hr light-dark cycles with free access to food and chow, and bred at The Center for Phenogenomics in Toronto ON, Canada. For all studies, mice of either sex were used at both the 3mo and 6mo timepoint. The specific ages of each animal for each experiment are documented in the respective results, method details, and/or figure legends of the study. A cohort of Ts65Dn mice, designated for the Ts65Dn+Fisetin condition, received a custom diet produced by Envigo, containing 500ppm Fisetin (Indofine Chemical, Cat: 528-48-3) in the chow, starting at 3mo of age, until euthanasia at 6mo. Quantitative PCR was performed on genomic DNA extracted from tail tips of all mice to confirm individual genotype. All genotyping primers used are listed in the key resources table.

### Tissue preparation for immunostaining

For immunostaining, Ts65Dn, Fisetin-treated (Ts65Dn+Fisetin), and euploid control (CTL) mice were perfused intra-aortically with a solution of 4% paraformaldehyde (PFA) while under isoflurane anesthesia at 3mo and 6mo (for untreated Ts65Dn and CTL groups at 3mo: *n* = 4 per condition; for Ts65Dn, Ts65Dn+Fisetin, and CTL groups at 6mo: *n = 3* per condition). Brains were removed and postfixed in 4% PFA at 4°C overnight, then washed 3× for 20 min in 1X PBS prior to cryoprotection in a 30% sucrose solution at 4°C for 3-4 days. The tissue was embedded in optimal cutting temperature (O.C.T.) compound (Fisher Scientific) and allowed to solidify at -80°C. Brains were then sliced coronally at 20µm on a cryostat and mounted onto 0.5% gelatin-coated superfrost slides (Fisher Scientific) and stored at -80°C long-term; three sequential sections were collected per slide.

### Senescence-associated-**β**-galactosidase (SA-**β**-gal) staining

SA-β-gal activity was determined on coronal brain sections using a SA-β-gal kit (Cell Signaling) according to the manufacturer’s instructions. Briefly, sections were incubated at 37°C for 20 min, washed 3× in 1X PBS for 10 min each, fixed for 10 min at room temperature using the kit’s fixative agent, and stained overnight for 16 hr using the SA-β-gal staining solution (pH 5.9-6.1, prepared according to kit instructions) in a sealed container placed in a CO_2_-free incubator. The sections were then washed in 1X PBS for 10 min and double-distilled water (DDW) for 5 min prior to coverslip mounting with mounting medium (ThermoFisher).

### Immunohistochemistry (IHC)

When coupled with SA-β-gal staining, the above protocol was followed with several additional steps. After washing in 1X PBS and deionized water, slides were washed in TBS (0.1 M Tris-HCl pH 7.5, 0.15 M NaCl in DDW) for 10 min, permeabilized with TBS-Tx (0.4% Triton X-100 in TBS) 2× for 10 min each, and blocked in blocking buffer (5% BSA, 0.4% Triton X-100 in TBS) at room temperature for 1 hr. Primary antibody was diluted in blocking buffer and applied to the slide, incubated in a humidified chamber overnight at 4°C. Staining was subsequently performed using HRP/DAB Detection Kits (Abcam) according to the manufacturer’s instructions, with minor adaptations. Briefly, sections were washed 4× in PBS-T (0.2% Triton X-100 in 1X PBS), incubated with biotinylated goat anti-polyvalent for 10 min, washed 4× in PBS-T, incubated with streptavidin peroxidase for 10 min, washed 4× in PBS-T, incubated with DAB mixture (2% DAB chromogen in DAB substrate) for 8-10 min, washed 4× in 1X PBS, and mounted with mounting medium (ThermoFisher).

### Immunofluorescence (IF)

Sections were incubated at 37°C for 20 min, washed 3× in 1X PBS for 10 min each, and blocked in blocking buffer (10% BSA, 6% NDS, 0.3% Triton X-100 in 1X PBS) at room temperature for 1 hr. Primary antibody was diluted in blocking buffer and applied to the slide, incubated in a humidified chamber overnight at 4°C. Sections were subsequently washed 3× in PBS-T (0.1% Triton X-100 in 1X PBS) for 10 min, and incubated with secondary antibody diluted in PBS-T for 1 hr at room temperature. After incubation, the sections were washed in 1X PBS and mounted with Fluoromount mounting medium (ThermoFisher).

### Antigen retrieval (AR)

When necessary, AR was coupled with the above IF protocol following the first round of 3× 10 min 1X PBS washes. Sodium citrate buffer (10 mM sodium citrate, 0.05% Tween-20, pH 6.0) was created according to Abcam protocol for heat-induced epitope retrieval. Slides were completely submerged in the buffer and heated for 20 min at full power in a domestic microwave once the solution reached boiling point. When 20 min elapsed, the heating vessel was removed and cooled in a bath of cold water until it reached room temperature. Slides were then washed 3× in 1X PBS for 10 min each and the aforementioned IF protocol was resumed beginning with 1 hr incubation in blocking buffer.

### Antibody concentrations

The primary antibodies used for immunostaining were goat anti-Pdgfrα (1:250, R&D Systems), rabbit anti-Pdgfrα (1:250, Abcam), rabbit anti-Olig2 (1:500, Abcam), mouse anti-CC1 (1:100, CalBioChem), rabbit anti-Iba1 (1:500, Wako), rabbit anti-LaminB1 (1:1000, Abcam; used with AR), mouse anti-NeuN (1:500, Millipore), mouse anti-Aqp4 (1:100, Abcam), rabbit anti-Ki67 (1:200, Abcam), rat anti-Cd68 (1:400, Abcam), rabbit anti-MBP (1:250, Abcam). The secondary antibodies used for immunostaining were Alexa Fluor 488-conjugated goat anti-rabbit IgG (1:500, ThermoFisher), Alexa Fluor 555- conjugated goat anti-mouse IgG (1:500, ThermoFisher), Alexa Fluor 555-conjugated goat anti-rat IgG (1:500, ThermoFisher), Alexa Fluor 488-conjugated donkey anti-goat (1:500, ThermoFisher), Alexa Fluor 555-conjugated donkey anti-mouse IgG (1:500, ThermoFisher), and Alexa Fluor 647-conjugated donkey anti-rabbit (1:500, ThermoFisher).

### Quantitative PCR (qPCR) analysis

qPCR was performed on 6mo tissue for Ts65Dn, Ts65Dn+Fisetin, and CTL groups at 6mo (*n = 4* per condition). mRNA expression levels of *Mbp* were measured by qPCR analysis. In brief, cortical tissue was lysed in 1 mL of Trizol (ThermoFisher) and total RNA was isolated according to the manufacturer’s guidelines. cDNA synthesis was performed using 2 μg of total RNA and the iScript cRNA synthesis kit (Bio-Rad). qPCR was performed using the Brilliant III ultra-Fast SYBR green qPCR master mix (Agilent Technologies) according to the manufacturer’s instructions. All primers used are listed in the key resources table. Data was normalized to *Gapdh* values.

### Slide Scanner imaging

Whole slide scans were acquired on a 3DHistech Pannoramic 250 Flash III SlideScanner using a Zeiss 40× 0.95 NA objective. The instrument was operated in extended focus mode (7 focal planes spanning 5 μm axial distance) to capture the entire cell volume across each tissue section. Maximum intensity projections of all the *z*-planes are shown in the specified figures. One image was captured per slide. The microscope is housed in the Imaging Facility at The Hospital for Sick Children in Toronto ON, Canada. Digital images obtained from the Slide Scanner were imported to the HALO Image Analysis Platform (Indica Labs) for subsequent analysis.

### Epifluorescence imaging

Fluorescence images were captured with a 40× water-immersion or 63× oil-immersion objective on a Zeiss Axio Imager.M2 upright microscope. Images were evenly captured across layers II-VI of the motor and somatosensory cortex on each tissue sample, for a total of 18 images per mouse. All imaging was captured using the Zen Blue software (Carl Zeiss Meditec). Digital images obtained from the Zeiss Axio Imager.M2 were then imported to the Volocity 3D Image Analysis Software (PerkinElmer) for subsequent analysis.

### Confocal imaging

High magnification brightfield images for Olig2/SA-β-gal and Pdgfrα/SA-β-gal analysis were acquired using a 40×/1.3 objective on Nikon Eclipse Ti2-E equipped with a Nikon DS10 color camera, running NIS Elements acquisition software. The microscope is housed in the Imaging Facility at The Hospital for Sick Children in Toronto ON, Canada.

### Tissue preparation for single-nucleus sequencing

For single-nucleus sequencing, cortical tissue from 6mo Ts65Dn and CTL (*n* = 3 per condition) mice were microdissected and snap frozen in liquid nitrogen. Frozen tissue was thawed in 1 ml Buffer HB (0.25 M sucrose, 25 mM KCl, 5 mM MgCl_2_, 20 mM Tricine-KOH pH 7.8, 0.15 mM spermine tetrahydrochloride, 0.5 mM spermidine trihydrochloride, 1 mM DTT). The tissue was transferred to a 7 mL dounce. 340 μL 5% IGEPAL CA-630 (Sigma-Aldrich) and 4 mL HB were added to the tissue and the tissue was homogenized with a tight pestle 10-15 times. The sample was transferred to a 15 mL tube and total solution brought to 10 mL with 50% iodixanol. 1 mL 30% iodixanol was layered on top of 1 mL 40% iodixanol in a separate corex tube (ThermoFisher). The 9 mL sample was layered on top of the iodixanol cushion. The sample was spun at 10,000× g for 18 min. 1 mL of sample at the 30-40% iodixanol interface was collected. After counting nuclei with a hemocytometer, the sample was diluted to 100,000 nuclei/mL with 30% iodixanol (with RNAsin) and subjected to single nuclear droplet encapsulation with the 10X Chromium platform (10X Genomics). Libraries were sequenced using the Illumina NovaSeq 6000 S4 platform at The Center for Applied Genomics (TCAG) at the Hospital for Sick Children in Toronto ON, Canada.

### Single-nucleus RNA sequencing (snRNA-seq) bioinformatics workflow

Raw reads were converted to FASTQs, mapped to the mm10 mouse reference genome (Ensembl 93), and demultiplexed to generate a per-cell count matrix using the CellRanger pipeline (10X Genomics). Cells from all snRNA-seq experiments were combined into a single Seurat object dataset and filtered through the Seurat pipeline using default parameters. Low quality cells with unique feature counts >2,500 or <200 and >5% mitochondrial counts were excluded. The data was then log-normalized and scaled by the default scale factor of 10,000. Variable features and principal components were then calculated using default values, returning 2,000 features for the dataset. A second round of scaling shifted the expression of each gene so that mean expression across cells was 0 and variance across cells was 1. Linear dimensional reduction was performed and the dimensionality of the dataset was determined. Clustering was performed by constructing a KNN graph and applying the Louvain algorithm. Dimensional reduction was performed with default values for uniform manifold approximation and projection (UMAP) and individual clusters were annotated based on expression of lineage-specific markers. The final snRNA-seq library contained 37,251 nuclei and represented all major cell types within the murine cortex. Differential gene expression (DGE) between the two conditions (Ts65Dn vs. CTL) for each cell type was assessed with the Seurat FindMarkers function using default MAST parameters and a log-fold change (logFC) threshold of 0.1. Benjamini-Hochberg corrected p-values were used for significance. Genes with FDR < 5% met statistical criteria for significant differential expression. Major cell-type annotations were assigned to clusters through manual inspection of canonical marker gene signals and confirmed through scPred’s reference-based mapping approach using the Allen Brain Atlas whole cortex and hippocampus 10X dataset^30^. Ligand-receptor (LR) analysis was performed on all identified cell clusters using CellChat with default parameters between the two conditions (Ts65Dn vs. CTL).

### Single-nucleus ATAC sequencing (snATAC-seq) bioinformatics workflow

Cells from all snATAC-seq experiments were combined into a single Seurat object dataset and filtered through the Signac pipeline using default parameters. Low quality cells were removed using filters for peak region fragments < 100,000, percent reads in peaks > 40, blacklist site ratio < 0.025, nucleosome signal < 4 and TSS enrichment score > 2. The data was then normalized via singular value decomposition (SVD) of the term frequency-inverse document frequency (TF-IDF) matrix and all identified variable features were used for linear dimensional reduction. Remaining pre-processing and dimensionality reduction was performed according to Signac recommendations using default parameters. A gene activity matrix was constructed by counting snATAC-seq peaks within the gene body and 2 kb upstream of the transcriptional start site (TSS) using protein-coding genes annotated in the Ensembl database. UMAP dimensionality reduction was performed with default values. The final snRNA-seq library contained 53,244 nuclei and represented all major cell types within the murine cortex. Differentially accessible regions (DARs) between the two conditions (Ts65Dn vs. CTL) for each cell type were calculated with the Signac FindMarkers function using default MAST parameters and a log-fold change (logFC) threshold of 0.1 and n_count_peaks as a latent variable. Benjamini-Hochberg corrected p-values were used for significance. Peaks with FDR < 5% met statistical criteria for significant differential accessibility. Major cell-type annotations were assigned to clusters through manual inspection of canonical marker gene signals and confirmed through label transfer using Seurat’s multimodal snRNA-seq to snATAC-seq integration pipeline. Integration showed 100% cluster label concordance between the two modalities. Transcription factor (TF) motif activity was estimated using the RunChromVAR wrapper in Signac. The positional weight matrix (PWM) was obtained from the JASPAR2020 database^145^. Differential motif activity was assessed using log_2_fold change (log_2_FC) threshold of 0.1 with a 0 pseudocount. Motifs with FDR < 5% met statistical criteria for significant binding activity.

### Fear conditioning

Behavioral testing was performed at The Centre for Phenogenomics (TCP) in Toronto, ON Canada. Animals were left in their home cage inside the testing room, anteroom or on the day-rack at least 30 min prior to testing. Mice were then transferred into individual testing chambers with grid floors. After 120 sec without any stimulus (which gives the mouse time to get used to the surrounding) a 30 sec, 3600 kHz 95 dB audible tone (CS, conditioned stimulus) followed and was co-terminated with a 2 sec foot shock of 0.75 mA (US, unconditioned stimulus). Following the shock, mice spent another 30 sec in the chamber prior to removal. Concomitantly, the animals associated the background context cues with the CS (conditioning). Mice were then returned to their holding room where they spent 24 hr before being returned to the testing room for additional screening. They were placed back into the same testing chambers for 300 sec without changes to the context/interior and no stimuli applied. After conditioning, the CS or the spatial context elicited, in the absence of the US, a central state of fear that was expressed as reduced locomotor activity or total lack of movement (freezing). The time spent immobile was used as a measure of learning/memory performance. This phase was followed by a 2 hr break, which mice spent in their home cage before being returned to the testing chamber with a changed context. The grid floor was covered with a plastic sheet, the ceiling was changed from a square to triangular shape, the olfactory cues were changed, and the noise levels were reduced. Mice spent 180 sec in this novel environment before a 180 sec 3600 kHz 95 dB audible tone (no foot shock) was played again. Following this phase animals, were placed back into their home cage and the test was completed. Clidox 1:5:1 and 70% EtOH solutions were used to disinfect surfaces between testing mice from different cages.

### Quantification of SA-**β**-gal activity

Two regions of interest (ROIs) were drawn across layers II-VI of the motor and somatosensory cortex on each tissue sample, for a total of six ROIs of equivalent area per mouse. The Multiplex IHC (v3.1.4) algorithm on HALO was used for all cell type specific SA-β-gal activity quantification and was optimized to detect cell type specific positivity based on the respective DAB staining. An ROI of approximately 3mm^2^ spanning the motor and somatosensory cortical regions was drawn for quantification of cell counts in the cortex across all biological replicates. A cytoplasmic radius was then measured and the percentage of cells that exhibited SA-β-gal positivity within the nuclear or associated cytoplasmic compartments was calculated.

### Quantification of Olig2/Pdgfr**α**/CC1 and Pdgfr**α**/Ki67 co-localization

The HighPlex FL (v4.1.3) algorithm on HALO was used for quantification of Olig2/Pdgfrα/CC1 and Pdgfrα/Ki67 colocalization positivity. The module was optimized to detect Olig2 or Pdgfrα-positive nuclear positivity for quantification of Olig2/Pdgfrα/CC1 and Pdgfrα/Ki67, respectively. A cytoplasmic radius was measured and the percentage of cells that exhibited concomitant nuclear or cytoplasmic compartment positivity for other markers was calculated accordingly. Thresholds for nuclear and cytoplasmic positivity were calibrated for each marker set and slide. An ROI of approximately 3mm^2^ spanning the motor and somatosensory cortical regions was drawn for quantification of cell counts in the cortex; an ROI of approximately 1mm^2^ was drawn medially along the CC for quantification of cell counts in the CC for all biological replicates.

### Quantification of nuclear Lmnb1 fluorescence intensity

Each image contained at least one Pdgfrα-positive cell. Pdgfrα-positive cells were manually selected and individually assessed for nuclear Lmnb1 colocalization within the imaged plane. The threshold for positive Lmnb1 staining was set to approximately one standard deviation (SD) from the mean across all sampled images. Normalized Lmnb1 intensity was then calculated for each Pdgfrα-positive cell by dividing the obtained Lmnb1 intensity value by the total surface area of the associated cell. A total of 100 Pdgfrα-positive cells were imaged from Ts65Dn, Ts65Dn+Fisetin, or CTL mice (*n* = 3 per condition) across the motor and somatosensory cortex.

### Quantification of Mbp fluorescence intensity

Myelin intensity thresholds were established based on parameters that allowed the maximum capture of Mbp stain across all images for all sampled conditions. Normalized Mbp intensity was then calculated for each image by dividing the obtained Mbp intensity value by the total surface area of all myelin sheaths within the imaged frame from images captures across the motor and somatosensory cortex..

### Immunostaining statistical analyses

Statistical significance was evaluated using a two-tailed student’s t-test for all experiments evaluating comparisons between Ts65Dn and CTL mice, or using an one-way ANOVA followed by a post-hoc Tukey test for all experiments evaluating comparisons between Ts65Dn, Ts65Dn+Fisetin, and CTL conditions. The statistical analyses were performed on R (The R Project for Statistical Computing). Error bars denote the standard deviation (SD) from the mean.

### Differential gene expression and differential accessibility statistical analysis

Statistics used to test differential gene expression and gene accessibility in the snRNA-seq and snATAC-seq data, respectively, were performed using the Seurat FindMarkers function. Genes or peaks with an adjusted p-value < 5% were considered statistically significant (Benjamini-Hochberg).

### GSEA and GO statistical analysis

False discovery rates reported in the GSEA and GO analyses were calculated using the Benjamini Hochberg method. FDR values < 5% were considered to be statistically significant.

### Quantification of senescence-associated gene signature

To perform transcriptomic gene set enrichment analysis for the senescence-associated gene signature, we used the previously published SenMayo^64^ published gene set and computed a normalized enrichment score (NES) for each cell type through the gProfiler package in R. Statistical significance was determined by cell types showing enrichment with FDR < 5%.

### Replicates

All immunostaining analyses were performed on at least 3 different brains each. All single-nucleus sequencing analyses were performed on 3 different brains each. Additional details of the statistical analyses can be found in the detailed descriptions of the immunostaining and computational methods above and, where relevant, in the results and the figure legends. For all analyses, *n* refers to number of animals analyzed.

## Data availability

Sequencing data have been deposited in GEO under accession code GSE225554. All data generated or analysed during this study are included in the manuscript and supporting files.

## Acknowledgements

We would like to thank Paul Paroutis and Kimberly Lau from the The Imaging Facility at the Hospital for Sick Children, Toronto ON, Canada for their assistance in setting up imaging protocols and analysis pipelines. We would like to thank Lois Kelsey, Ann Flenniken, and Igor Vukobradovic from The Centre for Phenogenomics Behavioral Phenotyping Core, Toronto ON, Canada for assisting with mouse behavioral analysis.

## Ethics

All animal use was approved by the Animal Care Committee of the Hospital for Sick Children in accordance with the Canadian Council of Animal Care policies (AUP #25-0374H).

## Supplemental figures

**Figure 1–figure supplement 1 A-D:**
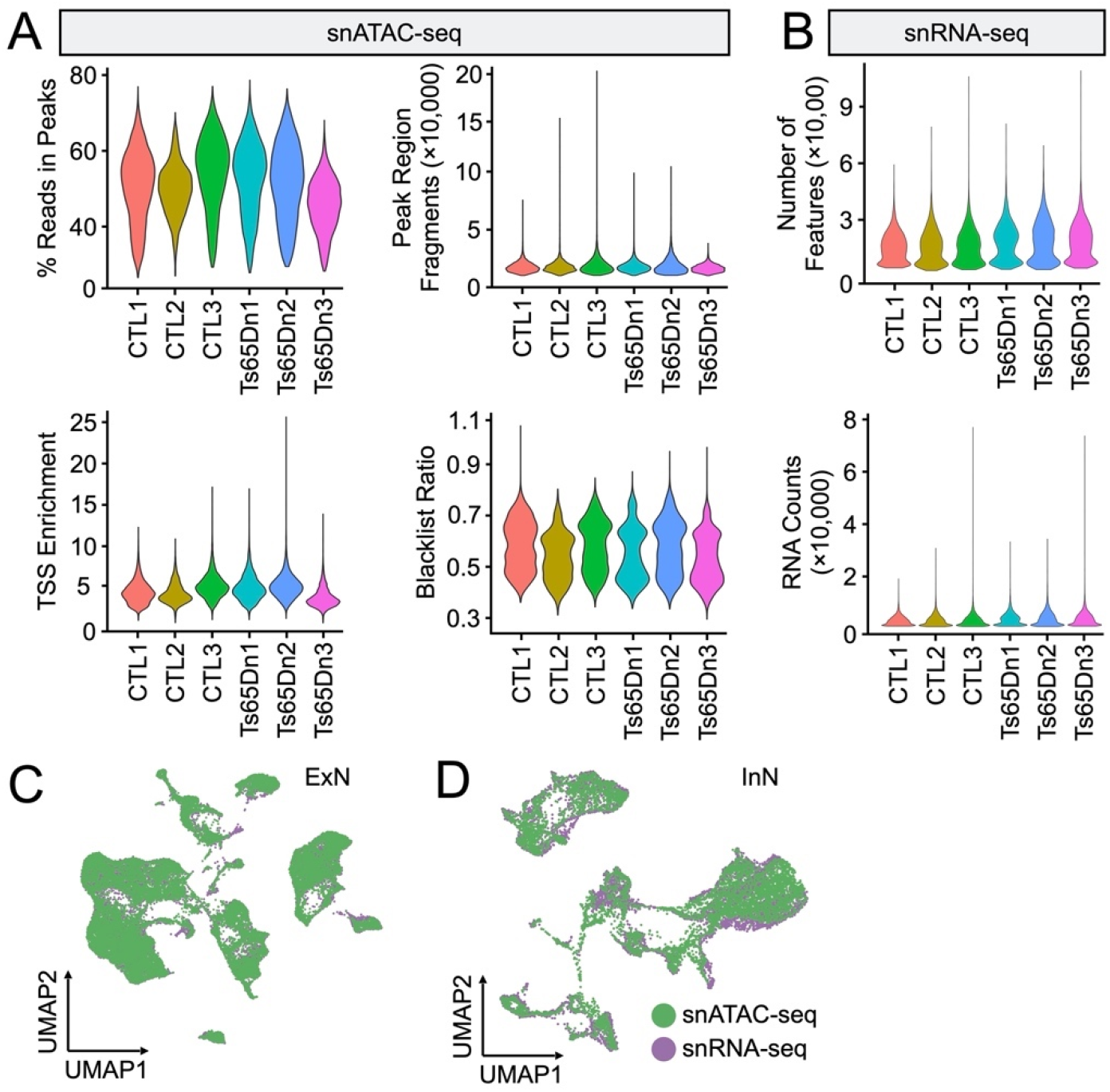
Quality control metric for multi-modal single-nucleus data and integration of snATAC- and snRNA-seq data for neuronal subsets, related to Figure 1. (A) quality control (QC) metrics for snATAC-seq data, including the percentage of reads in peaks, the number of peak region fragments, transcription start site (TSS) enrichment, blacklist ratio, and nucleosome signal per replicate from 6mo Ts65Dn and euploid control (CTL) mice. *n* = 3 biological replicates per condition. (B) as in (A), but depicting QC metrics for snRNA-seq data, including the number of features, total transcript counts, and percent mitochondrial content per replicate. (C) UMAP visualization of multi-omic integration of snATAC- and snRNA-seq datasets colored by originating dataset for the excitatory neuron (ExN) subset. (D) as in (C), but depicting multi-omic integration for the inhibitory neuron (InN) subset.

**Figure 6–figure supplement 1 A-D:**
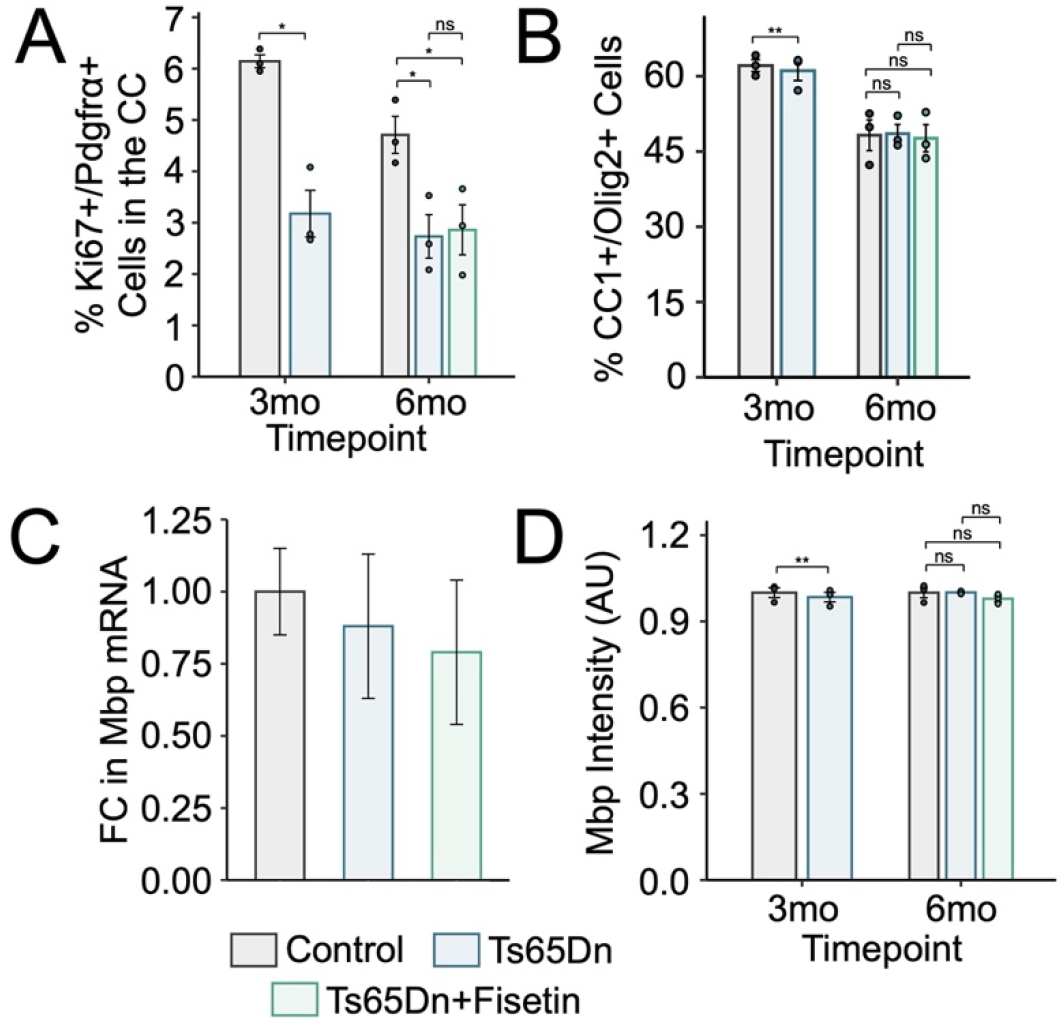
Quantification of cortical Mbp intensity, *Mbp* transcript-level, mOLs in the cortex, and proliferating OPCs in the CC, related to Figure 6. (A) Barplot of the percentage of Ki67+/ Pdgfrα+ cells (OPCs) in the CC, with a significant decrease in the number of OPCS in Ts65Dn at 3mo and 6mo, but no rescue in cell counts in 6mo Ts65Dn mice treated with Fisetin. Bars represent the average values for each condition (*n* = 3 mice per condition, *n* = 3 replicates per mouse), and dots represent the average values for each mouse per condition. Significance is determined using the two-tailed Student’s t-test at 3mo, or using the ANOVA test at 6mo. *p-value < 0.05; **p-value < 0.01, ***p-value < 0.001, ns: not significant; error bars represent the average ± 1 standard deviation (SD). (B) as in (A) but depicting the percentage of CC1+/Olig2+ cells (mOLs) in the cortex, with no significant difference in counts in any condition at eithet timepoint. (C) Barplot of the fold change (FC) in *Mbp* mRNA as measured through quantitative PCR (qPCR). Values were first normalized to *Gapdh*, and then normalized to the mean expression found in CTL replicates. Bars represent the average values for each condition (*n* = 3 mice per condition at 6mo, *n* = 4 replicates per mouse); error bars represent the average ± 1 standard deviation (SD). (D) as in (B), but depicting Mbp intensity in arbitrary units (AU). Values have been normalized to the mean intensity found in CTL replicates.

## References

1. Alexander, M., Petri, H., Ding, Y., Wandel, C., Khwaja, O., and Foskett, N. (2016). Morbidity and Medication in a Large Population of Individuals with Down Syndrome Compared to the General Population. Dev. Med. Child Neurol. 58, 246–254. 10.1111/dmcn.12868.

2. Kazemi, M., Salehi, M., and Kheirollahi, M. (2016). Down Syndrome: Current Status, Challenges and Future Perspectives. Int. J. Mol. Cell. Med. 5, 125–133.

3. Antonarakis, S.E., Skotko, B.G., Rafii, M.S., Strydom, A., Pape, S.E., Bianchi, D.W., Sherman, S.L., and Reeves, R.H. (2020). Down Syndrome. Nat. Rev. Dis. Primer 6, 9. 10.1038/s41572-019-0143-7.

4. Zigman, W.B., Devenny, D.A., KrinskyLMcHale, S.J., Jenkins, E.C., Urv, T.K., Wegiel, J., Schupf, N., and Silverman, W. (2008). Alzheimer’s Disease in Adults with Down Syndrome. In International Review of Research in Mental Retardation (Elsevier), pp. 103–145. 10.1016/S0074-7750(08)00004-9.

5. Sturgeon, X., and Gardiner, K.J. (2011). Transcript Catalogs of Human Chromosome 21 and Orthologous Chimpanzee and Mouse Regions. Mamm. Genome 22, 261–271. 10.1007/s00335-011-9321-y.

6. Becker, L., Mito, T., Takashima, S., and Onodera, K. (1991). Growth and Development of the Brain in Down Syndrome. Prog. Clin. Biol. Res. 373, 133–152.

7. Olmos-Serrano, J.L., Kang, H.J., Tyler, W.A., Silbereis, J.C., Cheng, F., Zhu, Y., Pletikos, M., Jankovic-Rapan, L., Cramer, N.P., Galdzicki, Z., et al. (2016). Down Syndrome Developmental Brain Transcriptome Reveals Defective Oligodendrocyte Differentiation and Myelination. Neuron 89, 1208–1222. 10.1016/j.neuron.2016.01.042.

8. Chakrabarti, L., Best, T.K., Cramer, N.P., Carney, R.S.E., Isaac, J.T.R., Galdzicki, Z., and Haydar, T.F. (2010). Olig1 and Olig2 Triplication Causes Developmental Brain Defects in Down Syndrome. Nat. Neurosci. 13, 927–934. 10.1038/nn.2600.

9. Barlow, G.M., Lyons, G.E., Richardson, J.A., Sarnat, H.B., and Korenberg, J.R. (2002). DSCAM: an Endogenous Promoter Drives Expression in the Developing CNS and Neural Crest. Biochem. Biophys. Res. Commun. 299, 1–6. 10.1016/S0006-291X(02)02548-2.

10. Chakrabarti, L., Galdzicki, Z., and Haydar, T.F. (2007). Defects in Embryonic Neurogenesis and Initial Synapse Formation in the Forebrain of the Ts65Dn Mouse Model of Down Syndrome. J. Neurosci. Off. J. Soc. Neurosci. 27, 11483–11495. 10.1523/JNEUROSCI.3406-07.2007.

11. Godfrey, M., and Lee, N.R. (2018). Memory Profiles in Down Syndrome Across Development: a Review of Memory Abilities Through the Lifespan. J. Neurodev. Disord. 10, 5. 10.1186/s11689-017-9220-y.

12. Teller, J.K., Russo, C., Debusk, L.M., Angelini, G., Zaccheo, D., Dagna-Bricarelli, F., Scartezzini, P., Bertolini, S., Mann, D.M.A., Tabaton, M., et al. (1996). Presence of Soluble Amyloid Β-Peptide Precedes Amyloid Plaque Formation in Down’s Syndrome. Nat. Med. 2, 93–95. 10.1038/nm0196-93.

13. Head, E., Lott, I.T., Wilcock, D.M., and Lemere, C.A. (2016). Aging in Down Syndrome and the Development of Alzheimer’s Disease Neuropathology. Curr. Alzheimer Res. 13, 18–29. 10.2174/1567205012666151020114607.

14. Flores-Aguilar, L., Iulita, M.F., Kovecses, O., Torres, M.D., Levi, S.M., Zhang, Y., Askenazi, M., Wisniewski, T., Busciglio, J., and Cuello, A.C. (2020). Evolution of Neuroinflammation Across the Lifespan of Individuals with Down Syndrome. Brain J. Neurol. 143, 3653–3671. 10.1093/brain/awaa326.

15. Wilcock, D.M. (2012). Neuroinflammation in the Aging Down Syndrome Brain; Lessons from Alzheimer’s Disease. Curr. Gerontol. Geriatr. Res. 2012, 170276. 10.1155/2012/170276.

16. Sullivan, K.D., Lewis, H.C., Hill, A.A., Pandey, A., Jackson, L.P., Cabral, J.M., Smith, K.P., Liggett, L.A., Gomez, E.B., Galbraith, M.D., et al. (2016). Trisomy 21 Consistently Activates the Interferon Response. eLife 5, e16220. 10.7554/eLife.16220.

17. Kong, X.-F., Worley, L., Rinchai, D., Bondet, V., Jithesh, P.V., Goulet, M., Nonnotte, E., Rebillat, A.S., Conte, M., Mircher, C., et al. (2020). Three Copies of Four Interferon Receptor Genes Underlie a Mild Type I Interferonopathy in Down Syndrome. J. Clin. Immunol. 40, 807–819. 10.1007/s10875-020-00803-9.

18. Lu, J., Esposito, G., Scuderi, C., Steardo, L., Delli-Bovi, L.C., Hecht, J.L., Dickinson, B.C., Chang, C.J., Mori, T., and Sheen, V. (2011). S100B and APP Promote a Gliocentric Shift and Impaired Neurogenesis in Down Syndrome Neural Progenitors. PloS One 6, e22126. 10.1371/journal.pone.0022126.

19. Wilcock, D.M., and Griffin, W.S.T. (2013). Down’s Syndrome, Neuroinflammation, and Alzheimer Neuropathogenesis. J. Neuroinflammation 10, 864. 10.1186/1742-2094-10-84.

20. Pinto, B., Morelli, G., Rastogi, M., Savardi, A., Fumagalli, A., Petretto, A., Bartolucci, M., Varea, E., Catelani, T., Contestabile, A., et al. (2020). Rescuing Over-activated Microglia Restores Cognitive Performance in Juvenile Animals of the Dp(16) Mouse Model of Down Syndrome. Neuron 108, 887–904.e12. 10.1016/j.neuron.2020.09.010.

21. Gupta, M., Dhanasekaran, A.R., and Gardiner, K.J. (2016). Mouse Models of Down Syndrome: Gene Content and Consequences. Mamm. Genome Off. J. Int. Mamm. Genome Soc. 27, 538–555. 10.1007/s00335-016-9661-8.

22. Reeves, R.H., Irving, N.G., Moran, T.H., Wohn, A., Kitt, C., Sisodia, S.S., Schmidt, C., Bronson, R.T., and Davisson, M.T. (1995). A Mouse Model for Down Syndrome Exhibits Learning and Behaviour Deficits. Nat. Genet. 11, 177–184. 10.1038/ng1095-177.

23. Hyde, L.A., Frisone, D.F., and Crnic, L.S. (2001). Ts65Dn Mice, A Model for Down Syndrome, Have Deficits in Context Discrimination Learning Suggesting Impaired Hippocampal Function. Behav. Brain Res. 118, 53–60. 10.1016/S0166-4328(00)00313-2.

24. Granholm, A.-C.E., Sanders, L.A., and Crnic, L.S. (2000). Loss of Cholinergic Phenotype in Basal Forebrain Coincides with Cognitive Decline in a Mouse Model of Down’s Syndrome. Exp. Neurol. 161, 647–663. 10.1006/exnr.1999.7289.

25. Satpathy, A.T., Granja, J.M., Yost, K.E., Qi, Y., Meschi, F., McDermott, G.P., Olsen, B.N., Mumbach, M.R., Pierce, S.E., Corces, M.R., et al. (2019). Massively Parallel Single-Cell Chromatin Landscapes of Human Immune Cell Development and Intratumoral T Cell Exhaustion. Nat. Biotechnol. 37, 925–936. 10.1038/s41587-019-0206-z.

26. Zheng, G.X.Y., Terry, J.M., Belgrader, P., Ryvkin, P., Bent, Z.W., Wilson, R., Ziraldo, S.B., Wheeler, T.D., McDermott, G.P., Zhu, J., et al. (2017). Massively Parallel Digital Transcriptional Profiling of Single Cells. Nat. Commun. 8, 14049. 10.1038/ncomms14049.

27. Shipp, S. (2007). Structure and Function of the Cerebral Cortex. Curr. Biol. 17, R443–R449. 10.1016/j.cub.2007.03.044.

28. Lee, N.R., Adeyemi, E.I., Lin, A., Clasen, L.S., Lalonde, F.M., Condon, E., Driver, D.I., Shaw, P., Gogtay, N., Raznahan, A., et al. (2016). Dissociations in Cortical Morphometry in Youth with Down Syndrome: Evidence for Reduced Surface Area but Increased Thickness. Cereb. Cortex 26, 2982–2990. 10.1093/cercor/bhv107.

29. Utagawa, E.C., Moreno, D.G., Schafernak, K.T., Arva, N.C., Malek-Ahmadi, M.H., Mufson, E.J., and Perez, S.E. (2022). Neurogenesis and Neuronal Differentiation in the Postnatal Frontal Cortex in Down Syndrome. Acta Neuropathol. Commun. 10, 86. 10.1186/s40478-022-01385-w.

30. Hao, Y., Hao, S., Andersen-Nissen, E., Mauck, W.M., Zheng, S., Butler, A., Lee, M.J., Wilk, A.J., Darby, C., Zager, M., et al. (2021). Integrated Analysis of Multimodal Single-Cell Data. Cell 184, 3573–3587.e29. 10.1016/j.cell.2021.04.048.

31. Stuart, T., Srivastava, A., Madad, S., Lareau, C.A., and Satija, R. (2021). Single-Cell Chromatin State Analysis with Signac. Nat. Methods 18, 1333–1341. 10.1038/s41592-021-01282-5.

32. Alquicira-Hernandez, J., Sathe, A., Ji, H.P., Nguyen, Q., and Powell, J.E. (2019). scPred: Accurate Supervised Method for Cell-Type Classification from Single-Cell RNA-Seq Data. Genome Biol. 20, 264. 10.1186/s13059-019-1862-5.

33. Yao, Z., van Velthoven, C.T.J., Nguyen, T.N., Goldy, J., Sedeno-Cortes, A.E., Baftizadeh, F., Bertagnolli, D., Casper, T., Chiang, M., Crichton, K., et al. (2021). A Taxonomy of Transcriptomic Cell Types Across the Isocortex and Hippocampal Formation. Cell 184, 3222–3241.e26. 10.1016/j.cell.2021.04.021.

34. Keller, D., Erö, C., and Markram, H. (2018). Cell Densities in the Mouse Brain: A Systematic Review. Front. Neuroanat. 12, 83. 10.3389/fnana.2018.00083.

35. Kobayashi, H., Saragai, S., Naito, A., Ichio, K., Kawauchi, D., and Murakami, F. (2015). Calm1 Signaling Pathway is Essential for the Migration of Mouse Precerebellar Neurons. Dev. Camb. Engl. 142, 375–384. 10.1242/dev.112680.

36. Mukherjee, C., Kling, T., Russo, B., Miebach, K., Kess, E., Schifferer, M., Pedro, L.D., Weikert, U., Fard, M.K., Kannaiyan, N., et al. (2020). Oligodendrocytes Provide Antioxidant Defense Function for Neurons by Secreting Ferritin Heavy Chain. Cell Metab. 32, 259–272.e10. 10.1016/j.cmet.2020.05.019.

37. Chaplot, K., Jarvela, T.S., and Lindberg, I. (2020). Secreted Chaperones in Neurodegeneration. Front. Aging Neurosci. 12, 268. 10.3389/fnagi.2020.00268.

38. Mouton-Liger, F., Thomas, S., Rattenbach, R., Magnol, L., Larigaldie, V., Ledru, A., Herault, Y., Verney, C., and Créau, N. (2011). PCP4 (PEP19) Overexpression Induces Premature Neuronal Differentiation Associated with Ca2+/Calmodulin-Dependent Kinase II-δ Activation in Mouse Models of Down Syndrome. J. Comp. Neurol. 519, 2779–2802. 10.1002/cne.22651.

39. Mi, J., Yang, Y., Yao, H., Huan, Z., Xu, C., Ren, Z., Li, W., Tang, Y., Fu, R., and Ge, X. (2021). Inhibition of Heat Shock Protein Family A Member 8 Attenuates Spinal Cord Ischemia-Reperfusion Injury via Astrocyte NF-κB/NLRP3 Inflammasome Pathway: HSPA8 Inhibition Protects Spinal Ischemia-Reperfusion Injury. J. Neuroinflammation 18, 170. 10.1186/s12974-021-02220-0.

40. Zhou, Q., Wang, S., and Anderson, D.J. (2000). Identification of a Novel Family of Oligodendrocyte Lineage-Specific Basic Helix-Loop-Helix Transcription Factors. Neuron 25, 331–343. 10.1016/s0896-6273(00)80898-3.

41. Zhou, Q., and Anderson, D.J. (2002). The bHLH Transcription Factors OLIG2 And OLIG1 Couple Neuronal and Glial Subtype Specification. Cell 109, 61–73. 10.1016/s0092-8674(02)00677-3.

42. Xu, R., Brawner, A.T., Li, S., Liu, J.-J., Kim, H., Xue, H., Pang, Z.P., Kim, W.-Y., Hart, R.P., Liu, Y., et al. (2019). OLIG2 Drives Abnormal Neurodevelopmental Phenotypes in Human iPSC-Based Organoid and Chimeric Mouse Models of Down Syndrome. Cell Stem Cell 24, 908–926.e8.10.1016/j.stem.2019.04.014.

43. Berto, G., Camera, P., Fusco, C., Imarisio, S., Ambrogio, C., Chiarle, R., Silengo, L., and Di Cunto, F. (2007). The Down Syndrome Critical Region Protein TTC3 Inhibits Neuronal Differentiation via RhoA and Citron Kinase. J. Cell Sci. 120, 1859–1867. 10.1242/jcs.000703.

44. O’Brien, R.J., and Wong, P.C. (2011). Amyloid Precursor Protein Processing and Alzheimer’s Disease. Annu. Rev. Neurosci. 34, 185–204. 10.1146/annurev-neuro-061010-113613.

45. Ermak, G., and Davies, K.J.A. (2013). Chronic High Levels of the RCAN1-1 Protein may Promote Neurodegeneration and Alzheimer Disease. Free Radic. Biol. Med. 62, 47–51. 10.1016/j.freeradbiomed.2013.01.016.

46. Gillespie, M., Jassal, B., Stephan, R., Milacic, M., Rothfels, K., Senff-Ribeiro, A., Griss, J., Sevilla, C., Matthews, L., Gong, C., et al. (2022). The Reactome Pathway Knowledgebase 2022. Nucleic Acids Res. 50, D687–D692. 10.1093/nar/gkab1028.

47. Izzo, A., Mollo, N., Nitti, M., Paladino, S., Calì, G., Genesio, R., Bonfiglio, F., Cicatiello, R., Barbato, M., Sarnataro, V., et al. (2018). Mitochondrial Dysfunction in Down Syndrome: Molecular Mechanisms and Therapeutic Targets. Mol. Med. 24, 2. 10.1186/s10020-018-0004-y.

48. Aivazidis, S., Coughlan, C.M., Rauniyar, A.K., Jiang, H., Liggett, L.A., Maclean, K.N., Roede, J.R. (2017). The Burden of Trisomy 21 Disrupts the Proteostasis Network in Down Syndrome. PLOS ONE 12, e0176307. 10.1371/journal.pone.0176307.

49. Zhu, P.J., Khatiwada, S., Cui, Y., Reineke, L.C., Dooling, S.W., Kim, J.J., Li, W., Walter, P., and Costa-Mattioli, M. (2019). Activation of the ISR Mediates the Behavioral and Neurophysiological Abnormalities in Down Syndrome. Science 366, 843–849. 10.1126/science.aaw5185.

50. Gingras, A.C., Raught, B., and Sonenberg, N. (1999). eIF4 Initiation Factors: Effectors of mRNA Recruitment to Ribosomes and Regulators of Translation. Annu. Rev. Biochem. 68, 913–963. 10.1146/annurev.biochem.68.1.913.

51. Aitken, C.E., Beznosková, P., Vlčkova, V., Chiu, W.-L., Zhou, F., Valášek, L.S., Hinnebusch, A.G., and Lorsch, J.R. (2016). Eukaryotic Translation Initiation Factor 3 Plays Distinct Roles at the mRNA Entry and Exit Channels of the Ribosomal Preinitiation Complex. eLife 5, e20934. 10.7554/eLife.20934.

52. Israel, A. (2010). The IKK Complex, a Central Regulator of NF- B Activation. Cold Spring Harb. Perspect. Biol. 2, a000158–a000158. 10.1101/cshperspect.a000158.

53. Renault-Mihara, F., and Okano, H. (2017). STAT3-Regulated RhoA Drives Reactive Astrocyte Dynamics. Cell Cycle 16, 1995–1996. 10.1080/15384101.2017.1377032.

54. Hazrati, A., Soudi, S., Malekpour, K., Mahmoudi, M., Rahimi, A., Hashemi, S.M., and Varma, R.S. (2022). Immune Cells-Derived Exosomes Function as a Double-Edged Sword: Role in Disease Progression and their Therapeutic Applications. Biomark. Res. 10, 30. 10.1186/s40364-022-00374-4.

55. Pabois, A., Devallière, J., Quillard, T., Coulon, F., Gérard, N., Laboisse, C., Toquet, C., and Charreau, B. (2014). The Disintegrin and Metalloproteinase ADAM10 Mediates a Canonical Notch-Dependent Regulation of IL-6 Through Dll4 in Human Endothelial Cells. Biochem. Pharmacol. 91, 510–521. 10.1016/j.bcp.2014.08.007.

56. Schroder, K., Hertzog, P.J., Ravasi, T., and Hume, D.A. (2004). Interferon-γ: an Overview of Signals, Mechanisms and Functions. J. Leukoc. Biol. 75, 163–189. 10.1189/jlb.0603252.

57. Tanaka, K. (2009). The Proteasome: Overview of Structure and Functions. Proc. Jpn. Acad. Ser. B 85, 12–36. 10.2183/pjab.85.12.

58. Park, M.H., Jo, M., Kim, Y.R., Lee, C.-K., and Hong, J.T. (2016). Roles of Peroxiredoxins in Cancer, Neurodegenerative Diseases and Inflammatory Diseases. Pharmacol. Ther. 163, 1–23. 10.1016/j.pharmthera.2016.03.018.

59. Scheffler, J.M., Sparber, F., Tripp, C.H., Herrmann, C., Humenberger, A., Blitz, J., Romani, N., Stoitzner, P., and Huber, L.A. (2014). LAMTOR2 Regulates Dendritic Cell Homeostasis Through FLT3-Dependent mTOR Signalling. Nat. Commun. 5, 5138. 10.1038/ncomms6138.

60. Ponroy Bally, B., and Murai, K.K. (2021). Astrocytes in Down Syndrome Across the Lifespan. Front. Cell. Neurosci. 15, 702685. 10.3389/fncel.2021.702685.

61. Jørgensen, O.S., Brooksbank, B.W.L., and Balázs, R. (1990). Neuronal Plasticity and Astrocytic Reaction in Down Syndrome and Alzheimer Disease. J. Neurol. Sci. 98, 63–79. 10.1016/0022-510X(90)90182-M.

62. Chen, C., Jiang, P., Xue, H., Peterson, S.E., Tran, H.T., McCann, A.E., Parast, M.M., Li, S., Pleasure, D.E., Laurent, L.C., et al. (2014). Role of Astroglia in Down’s Syndrome Revealed by Patient-Derived Human-Induced Pluripotent Stem Cells. Nat. Commun. 5, 4430. 10.1038/ncomms5430.

63. Puente-Bedia, A., Berciano, M.T., Martínez-Cué, C., Lafarga, M., and Rueda, N. (2022). Oxidative-Stress-Associated Proteostasis Disturbances and Increased DNA Damage in the Hippocampal Granule Cells of the Ts65Dn Model of Down Syndrome. Antioxidants 11, 2438. 10.3390/antiox11122438.

64. Ahmed, Md.M., Johnson, N.R., Boyd, T.D., Coughlan, C., Chial, H.J., and Potter, H. (2021). Innate Immune System Activation and Neuroinflammation in Down Syndrome and Neurodegeneration: Therapeutic Targets or Partners? Front. Aging Neurosci. 13, 718426. 10.3389/fnagi.2021.718426.

65. Guedj, F., Pennings, J.L., Massingham, L.J., Wick, H.C., Siegel, A.E., Tantravahi, U., and Bianchi, D.W. (2016). An Integrated Human/Murine Transcriptome and Pathway Approach To Identify Prenatal Treatments For Down Syndrome. Sci. Rep. 6, 32353. 10.1038/srep32353.

66. Kumari, R., and Jat, P. (2021). Mechanisms of Cellular Senescence: Cell Cycle Arrest and Senescence Associated Secretory Phenotype. Front. Cell Dev. Biol. 9, 645593. 10.3389/fcell.2021.645593.

67. Di Micco, R., Krizhanovsky, V., Baker, D., and d’Adda di Fagagna, F. (2021). Cellular Senescence in Ageing: From Mechanisms to Therapeutic Opportunities. Nat. Rev. Mol. Cell Biol. 22, 75–95. 10.1038/s41580-020-00314-w.

68. Saul, D., Kosinsky, R.L., Atkinson, E.J., Doolittle, M.L., Zhang, X., LeBrasseur, N.K., Pignolo, R.J., Robbins, P.D., Niedernhofer, L.J., Ikeno, Y., et al. (2021). A New Gene Set Identifies Senescent Cells and Predicts Senescence-Associated Pathways Across Tissues (Cell Biology) 10.1101/2021.12.10.472095.

69. Ou, Z., Sun, Y., Lin, L., You, N., Liu, X., Li, H., Ma, Y., Cao, L., Han, Y., Liu, M., et al. (2016). Olig2-Targeted G-Protein-Coupled Receptor Gpr17 Regulates Oligodendrocyte Survival in Response to Lysolecithin-Induced Demyelination. J. Neurosci. 36, 10560–10573. 10.1523/JNEUROSCI.0898-16.2016.

70. Chen, Y., Wu, H., Wang, S., Koito, H., Li, J., Ye, F., Hoang, J., Escobar, S.S., Gow, A., Arnett, H.A., et al. (2009). The Oligodendrocyte-Specific G Protein–Coupled Receptor GPR17 is a Cell Intrinsic Timer of Myelination. Nat. Neurosci. 12, 1398–1406. 10.1038/nn.2410.

71. Kim, M.J., Kim, W.S., Kim, D.O., Byun, J.-E., Huy, H., Lee, S.Y., Song, H.Y., Park, Y.-J., Kim, T.-D., Yoon, S.R., et al. (2017). Macrophage Migration Inhibitory Factor Interacts with Thioredoxin Interacting Protein and Induces NF-κB Activity. Cell. Signal. 34, 110–120. 10.1016/j.cellsig.2017.03.007.

72. Salminen, A., and Kaarniranta, K. (2011). Control of p53 and NF-κB Signaling by WIP1 and MIF: Role in Cellular Senescence and Organismal Aging. Cell. Signal. 23, 747–752. 10.1016/j.cellsig.2010.10.012.

73. Cabral-Pacheco, G.A., Garza-Veloz, I., Castruita-De la Rosa, C., Ramirez-Acuña, J.M., Perez-Romero, B.A., Guerrero-Rodriguez, J.F., Martinez-Avila, N., and Martinez-Fierro, M.L. (2020). The Roles of Matrix Metalloproteinases and Their Inhibitors in Human Diseases. Int. J. Mol. Sci. 21, 9739. 10.3390/ijms21249739.

74. Rosenberg, G.A. (2002). Matrix Metalloproteinases in Neuroinflammation. Glia 39, 279–291. 10.1002/glia.10108.

75. Mijit, M., Caracciolo, V., Melillo, A., Amicarelli, F., and Giordano, A. (2020). Role of p53 in the Regulation of Cellular Senescence. Biomolecules 10, 420. 10.3390/biom10030420.

76. Coppé, J.-P., Desprez, P.-Y., Krtolica, A., and Campisi, J. (2010). The Senescence-Associated Secretory Phenotype: The Dark Side of Tumor Suppression. Annu. Rev. Pathol. Mech. Dis. 5, 99–118. 10.1146/annurev-pathol-121808-102144.

77. Zhao, H., Ji, Q., Wu, Z., Wang, S., Ren, J., Yan, K., Wang, Z., Hu, J., Chu, Q., Hu, H., et al. (2022). Destabilizing Heterochromatin by APOE Mediates Senescence. Nat. Aging 2, 303–316. 10.1038/s43587-022-00186-z.

78. Mercurio, L., Lulli, D., Mascia, F., Dellambra, E., Scarponi, C., Morelli, M., Valente, C., Carbone, M.L., Pallotta, S., Girolomoni, G., et al. (2020). Intracellular Insulin-Like Growth Factor Binding Protein 2 (IGFBP2) Contributes to the Senescence of Keratinocytes in Psoriasis by Stabilizing Cytoplasmic p21. Aging 12, 6823–6851. 10.18632/aging.103045.

79. Chen, H.-A., Ho, Y.-J., Mezzadra, R., Adrover, J.M., Smolkin, R., Zhu, C., Woess, K., Bernstein, N., Schmitt, G., Fong, L., et al. (2023). Senescence Rewires Microenvironment Sensing to Facilitate Antitumor Immunity. Cancer Discov. 13, 432–453. 10.1158/2159-8290.CD-22-0528.

80. Jin, S., Guerrero-Juarez, C.F., Zhang, L., Chang, I., Ramos, R., Kuan, C.-H., Myung, P., Plikus, M.V., and Nie, Q. (2021). Inference and Analysis of Cell-Cell Communication using CellChat. Nat. Commun. 12, 1088. 10.1038/s41467-021-21246-9.

81. Papadimitriou, E., Mourkogianni, E., Ntenekou, D., Christopoulou, M., Koutsioumpa, M., and Lamprou, M. (2022). On the Role of Pleiotrophin and its Receptors in Development and Angiogenesis. Int. J. Dev. Biol. 66, 115–124. 10.1387/ijdb.210122ep.

82. Wang, X. (2020). Pleiotrophin: Activity and Mechanism. In Advances in Clinical Chemistry (Elsevier), pp. 51–89. 10.1016/bs.acc.2020.02.003.

83. McClain, C.R., Sim, F.J., and Goldman, S.A. (2012). Pleiotrophin Suppression of Receptor Protein Tyrosine Phosphatase-β/ζ Maintains the Self-Renewal Competence of Fetal Human Oligodendrocyte Progenitor Cells. J. Neurosci. 32, 15066–15075. 10.1523/JNEUROSCI.1320-12.2012.

84. Harroch, S., Furtado, G.C., Brueck, W., Rosenbluth, J., Lafaille, J., Chao, M., Buxbaum, J.D., and Schlessinger, J. (2002). A Critical Role for the Protein Tyrosine Phosphatase Receptor Type Z in Functional Recovery from Demyelinating Lesions. Nat. Genet. 32, 411–414. 10.1038/ng1004.

85. Bowden, E.T., Stoica, G.E., and Wellstein, A. (2002). Anti-Apoptotic Signaling of Pleiotrophin Through its Receptor, Anaplastic Lymphoma Kinase. J. Biol. Chem. 277, 35862–35868. 10.1074/jbc.M203963200.

86. Koutsioumpa, M., Drosou, G., Mikelis, C., Theochari, K., Vourtsis, D., Katsoris, P., Giannopoulou, E., Courty, J., Petrou, C., Magafa, V., et al. (2012). Pleiotrophin Expression and Role in Physiological Angiogenesis in Vivo: Potential Involvement of Nucleolin. Vasc. Cell 4, 4. 10.1186/2045-824X-4-4.

87. Reynolds, L.E., Watson, A.R., Baker, M., Jones, T.A., D’Amico, G., Robinson, S.D., Joffre, C., Garrido-Urbani, S., Rodriguez-Manzaneque, J.C., Martino-Echarri, E., et al. (2010). Tumor Angiogenesis is Reduced in the Tc1 Mouse Model of Down Syndrome. Nature 465, 813–817. 10.1038/nature09106.

88. Mei, L., and Xiong, W.-C. (2008). Neuregulin 1 in Neural Development, Synaptic Plasticity and Schizophrenia. Nat. Rev. Neurosci. 9, 437–452. 10.1038/nrn2392.

89. Cahill, M.E., Jones, K.A., Rafalovich, I., Xie, Z., Barros, C.S., Müller, U., and Penzes, P. (2012). Control of Interneuron Dendritic Growth through NRG1/Erbb4-Mediated Kalirin-7 Disinhibition. Mol. Psychiatry 17, 1, 99–107. 10.1038/mp.2011.35.

90. Ting, A.K., Chen, Y., Wen, L., Yin, D.-M., Shen, C., Tao, Y., Liu, X., Xiong, W.-C., and Mei, L. (2011). Neuregulin 1 Promotes Excitatory Synapse Development and Function in GABAergic Interneurons. J. Neurosci. 31, 15–25. 10.1523/JNEUROSCI.2538-10.2011.

91. Brinkmann, B.G., Agarwal, A., Sereda, M.W., Garratt, A.N., Müller, T., Wende, H., Stassart, R.M., Nawaz, S., Humml, C., Velanac, V., et al. (2008). Neuregulin-1/ErbB Signaling Serves Distinct Functions in Myelination of the Peripheral and Central Nervous System. Neuron 59, 581–595. 10.1016/j.neuron.2008.06.028.

92. Mei, L., and Nave, K.-A. (2014). Neuregulin-ERBB Signaling in the Nervous System and Neuropsychiatric Diseases. Neuron 83, 27–49. 10.1016/j.neuron.2014.06.007.

93. Zhang, D., Sliwkowski, M.X., Mark, M., Frantz, G., Akita, R., Sun, Y., Hillan, K., Crowley, C., Brush, J., and Godowski, P.J. (1997). Neuregulin-3 (NRG3): a Novel Neural Tissue-Enriched Protein that Binds and Activates Erbb4. Proc. Natl. Acad. Sci. U. S. A. 94, 9562–9567. 10.1073/pnas.94.18.9562.

94. Avramopoulos, D. (2018). Neuregulin 3 and its Roles in Schizophrenia Risk and Presentation. Am. J. Med. Genet. Part B Neuropsychiatr. Genet. Off. Publ. Int. Soc. Psychiatr. Genet. 177, 257–266. 10.1002/ajmg.b.32552.

95. Wang, Q., Li, M., Wu, T., Zhan, L., Li, L., Chen, M., Xie, W., Xie, Z., Hu, E., Xu, S., et al. (2022). Exploring Epigenomic Datasets by ChIPseeker. Curr. Protoc. 2. 10.1002/cpz1.585.

96. Lu, J., Lian, G., Zhou, H., Esposito, G., Steardo, L., Delli-Bovi, L.C., Hecht, J.L., Lu, Q.R., and Sheen, V. (2012). OLIG2 Over-Expression Impairs Proliferation of Human Down Syndrome Neural Progenitors. Hum. Mol. Genet. 21, 2330–2340. 10.1093/hmg/dds052.

97. Zhang, K., Chen, S., Yang, Q., Guo, S., Chen, Q., Liu, Z., Li, L., Jiang, M., Li, H., Hu, J., et al. (2022). The Oligodendrocyte Transcription Factor 2 OLIG2 Regulates Transcriptional Repression During Myelinogenesis in Rodents. Nat. Commun. 13, 1423. 10.1038/s41467-022-29068-z.

98. Yeh, H., and Ikezu, T. (2019). Transcriptional and Epigenetic Regulation of Microglia in Health and Disease. Trends Mol. Med. 25, 96–111. 10.1016/j.molmed.2018.11.004.

99. Zusso, M., Methot, L., Lo, R., Greenhalgh, A.D., David, S., and Stifani, S. (2012). Regulation of Postnatal Forebrain Amoeboid Microglial Cell Proliferation and Development by the Transcription Factor Runx1. J. Neurosci. 32, 11285–11298. 10.1523/JNEUROSCI.6182-11.2012.

100. Choudhry, H., Aggarwal, M., and Pan, P.-Y. (2021). Mini-Review: Synaptojanin 1 and its Implications in Membrane Trafficking. Neurosci. Lett. 765, 136288. 10.1016/j.neulet.2021.136288.

101. Miranda, A.M., Herman, M., Cheng, R., Nahmani, E., Barrett, G., Micevska, E., Fontaine, G., Potier, M.-C., Head, E., Schmitt, F.A., et al. (2018). Excess Synaptojanin 1 Contributes to Place Cell Dysfunction and Memory Deficits in the Aging Hippocampus in Three Types of Alzheimer’s Disease. Cell Rep. 23, 2967–2975. 10.1016/j.celrep.2018.05.011.

102. Raudvere, U., Kolberg, L., Kuzmin, I., Arak, T., Adler, P., Peterson, H., and Vilo, J. (2019). gProfiler: a Web Server for Functional Enrichment Analysis and Conversions of Gene Lists (2019 Update). Nucleic Acids Res. 47, W191–W198. 10.1093/nar/gkz369.

103. Suhail, T.V., Singh, P., and Manna, T.K. (2015). Suppression of Centrosome Protein TACC3 Induces G1 Arrest and Cell Death through Activation Of p38-p53-p21 Stress Signaling Pathway. Eur. J. Cell Biol. 94, 90–100. 10.1016/j.ejcb.2014.12.001.

104. Lee, C.S., Bhaduri, A., Mah, A., Johnson, W.L., Ungewickell, A., Aros, C.J., Nguyen, C.B., Rios, E.J., Siprashvili, Z., Straight, A., et al. (2014). Recurrent Point Mutations in the Kinetochore Gene KNSTRN in Cutaneous Squamous Cell Carcinoma. Nat. Genet. 46, 1060–1062. 10.1038/ng.3091.

105. Cheung, A.K.L., Ip, J.C.Y., Chu, A.C.H., Cheng, Y., Leong, M.M.L., Ko, J.M.Y., Shuen, W.H., Lung, H.L., and Lung, M.L. (2015). PTPRG Suppresses Tumor Growth and Invasion via Inhibition of Akt Signaling in Nasopharyngeal Carcinoma. Oncotarget 6, 13434–13447. 10.18632/oncotarget.3876.

106. Jagadish, N., Gupta, N., Agarwal, S., Parashar, D., Sharma, A., Fatima, R., Topno, A.P., Kumar, V., and Suri, A. (2016). Sperm-Associated Antigen 9 (SPAG9) Promotes the Survival and Tumor Growth of Triple-Negative Breast Cancer Cells. Tumour Biol. J. Int. Soc. Oncodevelopmental Biol. Med. 37, 13101–13110. 10.1007/s13277-016-5240-6.

107. Jang, D.G., Sim, H.J., Song, E.K., Kwon, T., and Park, T.J. (2020). Extracellular Matrixes and Neuroinflammation. BMB Rep. 53, 491–499. 10.5483/BMBRep.2020.53.10.156.

108. Ghorbani, S., and Yong, V.W. (2021). The Extracellular Matrix as Modifier of Neuroinflammation and Remyelination in Multiple Sclerosis. Brain 144, 1958–1973. 10.1093/brain/awab059.

109. Tewari, B.P., Chaunsali, L., Prim, C.E., and Sontheimer, H. (2022). A Glial Perspective on the Extracellular Matrix and Perineuronal Net Remodeling in the Central Nervous System. Front. Cell. Neurosci. 16, 1022754. 10.3389/fncel.2022.1022754.

110. Nacarelli, T., Liu, P., and Zhang, R. (2017). Epigenetic Basis of Cellular Senescence and Its Implications in Aging. Genes 8, 343. 10.3390/genes8120343.

111. Sadaie, M., Salama, R., Carroll, T., Tomimatsu, K., Chandra, T., Young, A.R.J., Narita, M., Pérez-Mancera, P.A., Bennett, D.C., Chong, H., et al. (2013). Redistribution of the Lamin B1 Genomic Binding Profile Affects Rearrangement of Heterochromatic Domains and SAHF Formation During Senescence. Genes Dev. 27, 1800–1808. 10.1101/gad.217281.113.

112. Sidler, C., Kovalchuk, O., and Kovalchuk, I. (2017). Epigenetic Regulation of Cellular Senescence and Aging. Front. Genet. 8, 138. 10.3389/fgene.2017.00138.

113. Schep, A.N., Wu, B., Buenrostro, J.D., and Greenleaf, W.J. (2017). ChromVAR: Inferring Transcription-Factor-Associated Accessibility from Single-Cell Epigenomic Data. Nat. Methods 14, 975–978. 10.1038/nmeth.4401.

114. Mo, J., Kim, C.-H., Lee, D., Sun, W., Lee, H.W., and Kim, H. (2015). Early Growth Response 1 (Egr-1) Directly Regulates GABAA Receptor α2, α4, and θ Subunits in the Hippocampus. J. Neurochem. 133, 489–500. 10.1111/jnc.13077.

115. Bernhardt, C., Sock, E., Fröb, F., Hillgärtner, S., Nemer, M., and Wegner, M. (2022). KLF9 and KLF13 Transcription Factors Boost Myelin Gene Expression in Oligodendrocytes as Partners of SOX10 and MYRF. Nucleic Acids Res. 50, 11509–11528. 10.1093/nar/gkac953.

116. Liu, T., Zhang, L., Joo, D., and Sun, S.-C. (2017). NF-κB Signaling in Inflammation. Signal Transduct. Target. Ther. 2, 17023. 10.1038/sigtrans.2017.23.

117. Preciados, M., Yoo, C., and Roy, D. (2016). Estrogenic Endocrine Disrupting Chemicals Influencing NRF1 Regulated Gene Networks in the Development of Complex Human Brain Diseases. Int. J. Mol. Sci. 17, 2086. 10.3390/ijms17122086.

118. Morabito, S., Miyoshi, E., Michael, N., Shahin, S., Martini, A.C., Head, E., Silva, J., Leavy, K., Perez-Rosendahl, M., and Swarup, V. (2021). Single-Nucleus Chromatin Accessibility and Transcriptomic Characterization of Alzheimer’s Disease. Nat. Genet. 53, 1143–1155. 10.1038/s41588-021-00894-z.

119. Satoh, J., Kawana, N., and Yamamoto, Y. (2013). Pathway Analysis of ChIP-Seq-Based NRF1 Target Genes Suggests a Logical Hypothesis of their Involvement in the Pathogenesis of Neurodegenerative Diseases. Gene Regul. Syst. Biol. 7, GRSB.S13204. 10.4137/GRSB.S13204.

120. Ogata, T., Ueno, T., Hoshikawa, S., Ito, J., Okazaki, R., Hayakawa, K., Morioka, K., Yamamoto, S., Nakamura, K., Tanaka, S., et al. (2011). Hes1 Functions Downstream of Growth Factors to Maintain Oligodendrocyte Lineage Cells in the Early Progenitor Stage. Neuroscience 176, 132–141. 10.1016/j.neuroscience.2010.12.015.

121. Fan, H., Zhao, J., Yan, J., Du, G., Fu, Q., Shi, J., Yang, Y., Du, X., and Bai, X. (2018). Effect of Notch1 Gene on Remyelination in Multiple Sclerosis in Mouse Models of Acute Demyelination. J. Cell. Biochem. 119, 9284–9294. 10.1002/jcb.27197.

122. Hu, N., and Zou, L. (2022). Multiple Functions of Hes Genes in the Proliferation and Differentiation of Neural Stem Cells. Ann. Anat. Anat. Anz. 239, 151848. 10.1016/j.aanat.2021.151848.

123. Li, S., Guan, H., Zhang, Y., Li, S., Li, K., Hu, S., Zuo, E., Zhang, C., Zhang, X., Gong, G., et al. (2021). Bone Marrow Mesenchymal Stem Cells Promote Remyelination in Spinal Cord by Driving Oligodendrocyte Progenitor Cell Differentiation via TNFα/Relb-Hes1 Pathway: a Rat Model Study of 2,5-Hexanedione-Induced Neurotoxicity. Stem Cell Res. Ther. 12, 436. 10.1186/s13287-021-02518-z.

124. Freund, A., Laberge, R.-M., Demaria, M., and Campisi, J. (2012). Lamin B1 Loss is a Senescence-Associated Biomarker. Mol. Biol. Cell 23, 2066–2075. 10.1091/mbc.E11-10-0884.

125. Zhu, Y., Tchkonia, T., Pirtskhalava, T., Gower, A.C., Ding, H., Giorgadze, N., Palmer, A.K., Ikeno, Y., Hubbard, G.B., Lenburg, M., et al. (2015). The Achilles’ Heel of Senescent Cells: From Transcriptome to Senolytic Drugs. Aging Cell 14, 644–658. 10.1111/acel.12344.

126. Elsallabi, O., Patruno, A., Pesce, M., Cataldi, A., Carradori, S., and Gallorini, M. (2022). Fisetin as a Senotherapeutic Agent: Biopharmaceutical Properties and Crosstalk between Cell Senescence and Neuroprotection. Mol. Basel Switz. 27, 738. 10.3390/molecules27030738.

127. Camell, C.D., Yousefzadeh, M.J., Zhu, Y., Prata, L.G.P.L., Huggins, M.A., Pierson, M., Zhang, L., O’Kelly, R.D., Pirtskhalava, T., Xun, P., et al. (2021). Senolytics Reduce Coronavirus-Related Mortality in Old Mice. Science 373, eabe4832. 10.1126/science.abe4832.

128. Thomas, A.L., Lehn, M.A., Janssen, E.M., Hildeman, D.A., and Chougnet, C.A. (2022). Naturally Aged Microglia Exhibit Phagocytic Dysfunction Accompanied by Gene Expression Changes Reflective of Underlying Neurologic Disease. Sci. Rep. 12, 19471. 10.1038/s41598-022-21920-y.

129. Safaiyan, S., Kannaiyan, N., Snaidero, N., Brioschi, S., Biber, K., Yona, S., Edinger, A.L., Jung, S., Rossner, M.J., and Simons, M. (2016). Age-Related Myelin Degradation Burdens the Clearance Function of Microglia During Aging. Nat. Neurosci. 19, 995–998. 10.1038/nn.4325.

130. Marques, L., Johnson, A.A., and Stolzing, A. (2020). Doxorubicin Generates Senescent Microglia that Exhibit Altered Proteomes, Higher Levels of Cytokine Secretion, and a Decreased Ability to Internalize Amyloid β. Exp. Cell Res. 395, 112203. 10.1016/j.yexcr.2020.112203.

131. Stogsdill, J.A., Kim, K., Binan, L., Farhi, S.L., Levin, J.Z., and Arlotta, P. (2022). Pyramidal Neuron Subtype Diversity Governs Microglia States in the Neocortex. Nature 608, 750–756. 10.1038/s41586-022-05056-7.

132. Reiche, L., Küry, P., and Göttle, P. (2019). Aberrant Oligodendrogenesis in Down Syndrome: Shift in Gliogenesis? Cells 8, 1591. 10.3390/cells8121591.

133. Falcão, A.M., van Bruggen, D., Marques, S., Meijer, M., Jäkel, S., Agirre, E., Samudyata, Floriddia, E.M., Vanichkina, D.P., ffrench-Constant, C., et al. (2018). Disease-Specific Oligodendrocyte Lineage Cells Arise in Multiple Sclerosis. Nat. Med. 24, 1837–1844. 10.1038/s41591-018-0236-y.

134. Marisca, R., Hoche, T., Agirre, E., Hoodless, L.J., Barkey, W., Auer, F., Castelo-Branco, G., and Czopka, T. (2020). Functionally Distinct Subgroups of Oligodendrocyte Precursor Cells Integrate Neural Activity and Execute Myelin Formation. Nat. Neurosci. 23, 363–374. 10.1038/s41593-019-0581-2.

135. Neumann, B., Segel, M., Ghosh, T., Zhao, C., Tourlomousis, P., Young, A., Förster, S., Sharma, A., Chen, C.Z.-Y., Cubillos, J.F., et al. (2021). Myc Determines the Functional Age State of Oligodendrocyte Progenitor Cells. Nat. Aging 1, 826–837. 10.1038/s43587-021-00109-4.

136. Kirby, L., Jin, J., Cardona, J.G., Smith, M.D., Martin, K.A., Wang, J., Strasburger, H., Herbst, L., Alexis, M., Karnell, J., et al. (2019). Oligodendrocyte Precursor Cells Present Antigen and are Cytotoxic Targets in Inflammatory Demyelination. Nat. Commun. 10, 3887. 10.1038/s41467-019-11638-3.

137. Araya, P., Waugh, K.A., Sullivan, K.D., Núñez, N.G., Roselli, E., Smith, K.P., Granrath, R.E., Rachubinski, A.L., Enriquez Estrada, B., Butcher, E.T., et al. (2019). Trisomy 21 Dysregulates T Cell Lineages Toward an Autoimmunity-Prone State Associated with Interferon Hyperactivity. Proc. Natl. Acad. Sci. U. S. A. 116, 24231–24241. 10.1073/pnas.1908129116.

138. Waugh, K.A., Araya, P., Pandey, A., Jordan, K.R., Smith, K.P., Granrath, R.E., Khanal, S., Butcher, E.T., Estrada, B.E., Rachubinski, A.L., et al. (2019). Mass Cytometry Reveals Global Immune Remodeling with Multi-lineage Hypersensitivity to Type I Interferon in Down Syndrome. Cell Rep. 29, 1893–1908.e4. 10.1016/j.celrep.2019.10.038.

139. Jin, M., Xu, R., Wang, L., Alam, M.M., Ma, Z., Zhu, S., Martini, A.C., Jadali, A., Bernabucci, M., Xie, P., et al. (2022). Type-I-Interferon Signaling Drives Microglial Dysfunction and Senescence in Human IPSC Models of Down Syndrome and Alzheimer’s Disease. Cell Stem Cell 29, 1135–1153.e8. 10.1016/j.stem.2022.06.007.

140. Zhang, P., Kishimoto, Y., Grammatikakis, I., Gottimukkala, K., Cutler, R.G., Zhang, S., Abdelmohsen, K., Bohr, V.A., Misra Sen, J., Gorospe, M., et al. (2019). Senolytic Therapy Alleviates Aβ-Associated Oligodendrocyte Progenitor Cell Senescence and Cognitive Deficits in an Alzheimer’s Disease Model. Nat. Neurosci. 22, 719–728. 10.1038/s41593-019-0372-9.

141. Meharena, H.S., Marco, A., Dileep, V., Lockshin, E.R., Akatsu, G.Y., Mullahoo, J., Watson, L.A., Ko, T., Guerin, L.N., Abdurrob, F., et al. (2022). Down-Syndrome-Induced Senescence Disrupts the Nuclear Architecture of Neural Progenitors. Cell Stem Cell 29, 116–130.e7. 10.1016/j.stem.2021.12.002.

142. Choi, J.H.K., Berger, J.D., Mazzella, M.J., Morales-Corraliza, J., Cataldo, A.M., Nixon, R.A., Ginsberg, S.D., Levy, E., and Mathews, P.M. (2009). Age-Dependent Dysregulation of Brain Amyloid Precursor Protein in the Ts65Dn Down Syndrome Mouse Model. J. Neurochem. 110, 1818–1827. 10.1111/j.1471-4159.2009.06277.x.

143. Escorihuela, R.M., Fernández-Teruel, A., Vallina, I.F., Baamonde, C., Lumbreras, M.A., Dierssen, M., Tobeña, A., and Flórez, J. (1995). A Behavioral Assessment of Ts65Dn Mice: A Putative Down Syndrome Model. Neurosci. Lett. 199, 143–146. 10.1016/0304-3940(95)12052-6.

144. Duchon, A., del Mar Muñiz Moreno, M., Chevalier, C., Nalesso, V., Andre, P., Fructuoso-Castellar, M., Mondino, M., Po, C., Noblet, V., Birling, M.-C., et al. (2022). Ts66Yah, a Mouse Model of Down Syndrome with Improved Construct and Face Validity. Dis. Model. Mech. 15, dmm049721. 10.1242/dmm.049721.

145. Fornes, O., Castro-Mondragon, J.A., Khan, A., van der Lee, R., Zhang, X., Richmond, P.A., Modi, B.P., Correard, S., Gheorghe, M., Baranašić, D., et al. (2019). JASPAR 2020: Update of the Open-Access Database of Transcription Factor Binding Profiles. Nucleic Acids Res., gkz1001. 10.1093/nar/gkz1001.

